# Single Cell Transcriptomics Reveals Cell Type Specific Diversification in Human Heart Failure

**DOI:** 10.1101/2021.07.06.451312

**Authors:** Andrew L. Koenig, Irina Shchukina, Prabhakar S Andhey, Konstantin Zaitsev, Lulu Lai, Junedh Amrute, Geetika Bajpai, Andrea Bredemeyer, Gabriella Smith, Cameran Jones, Emily Terrebonne, Stacey L. Rentschler, Maxim N. Artyomov, Kory J. Lavine

## Abstract

Heart failure represents a major cause of morbidity and mortality worldwide. Single cell transcriptomics have revolutionized our understanding of cell composition and associated gene expression across human tissues. Through integrated analysis of single cell and single nucleus RNA sequencing data generated from 45 individuals, we define the cell composition of the healthy and failing human heart. We identify cell specific transcriptional signatures of heart failure and reveal the emergence of disease associated cell states. Intriguingly, cardiomyocytes converge towards a common disease associated cell state, while fibroblasts and myeloid cells undergo dramatic diversification. Endothelial cells and pericytes display global transcriptional shifts without changes in cell complexity. Collectively, our findings provide a comprehensive analysis of the cellular and transcriptomic landscape of human heart failure, identify cell type specific transcriptional programs and states associated with disease, and establish a valuable resource for the investigation of human heart failure.

## Introduction

Single cell and single nucleus RNA sequencing (scRNAseq, snRNAseq) represent powerful new tools to identify cell types and their respective transcriptional signatures that reside within healthy and diseased tissues. Prior to the development of these technologies, our understanding of the cells that comprise human tissues and organs was restricted to routine histology and immunostaining analyses performed many decades ago. The rapid deployment of single cell sequencing has revolutionized the field and resulted in the identification of previously unrecognized cell populations including disease specific cell states across a wide range of structures including brain, lung, liver, kidney, and various malignancies.^1–5^

Recently, scRNAseq and snRNAseq was performed on healthy human heart tissue^6,7^. These studies yielded new information pertaining to common and rare cell populations within the healthy heart. Cardiomyocytes, fibroblasts, endothelial cells, pericytes, smooth muscles cells, myeloid cells, lymphoid cells, adipocytes, and neuronal cells were readily identified and analyzed across anatomical sites. Distinct transcriptional states of atrial and ventricular cardiomyocytes were identified and validated using RNA *in situ* hybridization. Surprising diversity was also observed amongst perivascular and immune cell types including transcriptional signatures specific to different regions of heart. At present, little is understood regarding the functional relevance of cell diversity within major cardiac cell populations. Furthermore, the impact of cardiac disease on cell composition remains to be rigorously investigated.

Heart failure represents a major cause of morbidity and mortality worldwide and imparts large costs on health care systems.^8,9^ While bulk RNA sequencing has yielded important insights into disease mechanisms that contribute to heart failure pathogenesis^10^, cell specific information is lost and much remains to be learned regarding the roles of individual cell types. Identification of disease associated cell specific programs may provide the insights and opportunities necessary to develop new therapies for heart failure.

Herein, we performed snRNAseq and scRNAseq on a large cohort of heart specimens obtained from healthy subjects and chronic heart failure patients. We identified 15 major cardiac cell types from 45 individuals and explored the extent of cell diversity within each of these populations. Unsupervised clustering, differential gene expression, and trajectory analyses revealed cell type specific transcriptional programs and emergence of disease associated cell states in the context of heart failure. We observed only subtle differences related to age and sex among non-diseased donor samples. Our data provide a comprehensive analysis of the cellular and transcriptomic landscape of the healthy and failing human heart and will serve as a valuable resource to the scientific community.

## Results

### Single nucleus and single cell RNA sequencing reveals the cellular landscape of the human heart

To define the cellular and transcriptional landscape of the healthy and failing human heart, we obtained left ventricular (LV) cardiac tissue specimens from 28 non-diseased donors and 17 subjects with dilated (nonischemic) cardiomyopathy (DCM). Non-diseased tissues were acquired from prospective donor hearts with normal LV function that were not used for transplantation due to the lack of a suitable recipient. DCM tissue was obtained from subjects undergoing implantation of a left ventricular assist device or explanted hearts collected at the time of transplantation. Transmural myocardial samples from the apical and anterior segments of the LV were processed for either single nucleus RNA sequencing (snRNAseq, n=38) or single cell RNA sequencing (scRNAseq, n=7) using the 10x Genomics 5’ Single Cell platform (**Tables S1-S3, Fig. 1A**).

**Figure 1.**
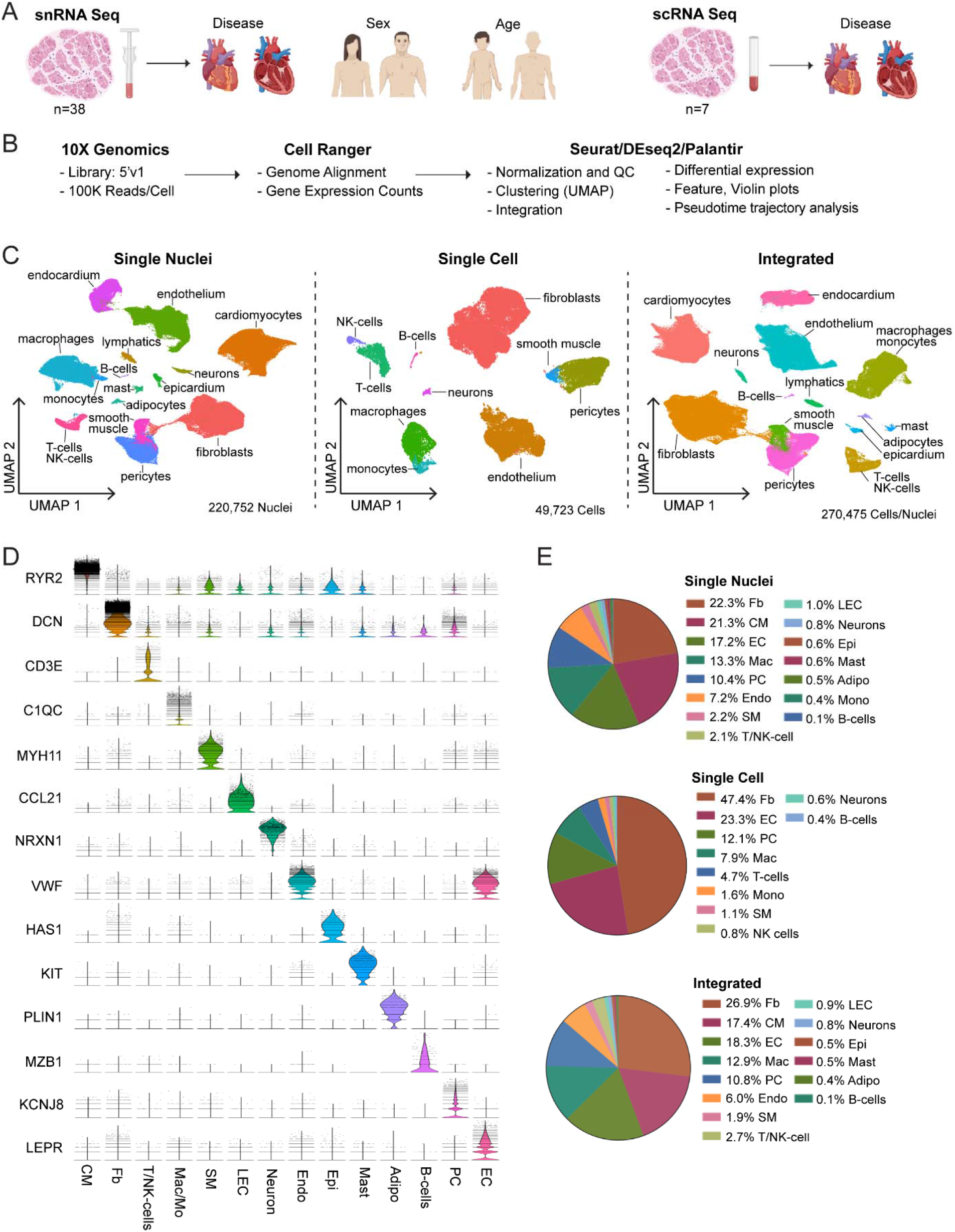
Cellular Composition of the Healthy and Failing Human Heart. **A,** Schematic depicting design of the single nucleus and single cell RNA sequencing experiments. Transmural sections were obtained from the apical anterior wall of the left ventricle during donor heart procurement, LVAD implantation, or heart transplantation for comparison of disease, sex, and age (single nucleus RNA sequencing: n=26 donor control, n=12 dilated cardiomyopathy; single cell RNA sequencing: n=2 donor control, n=5 dilated cardiomyopathy). **B,** The analysis pipeline included tissue processing and single cell barcoded library generation (10X genomics 5’v1 kit), sequence alignment (Cell Ranger), and further analysis using R and Python packages (Seurat, DEseq2, Palantir). **C,** Unsupervised UMAP clustering of 220,752 nuclei, 49,723 cells, and an integrated dataset combining single nucleus and single cell RNA sequencing data after QC and data filtering. **D,** Violin plots generated from the integrated dataset displaying characteristic marker genes of each identified cell population. **E,** Pie chart showing the proportion of cells within the single nucleus RNA sequencing, single cell RNA sequencing, and integrated datasets.

Single nucleus and single cell libraries were sequenced, aligned to the human reference genome, filtered for quality control, and unsupervised clustering, integration, and expression analysis performed using Seurat (**Fig. 1B, Fig. S1, Fig. S2**). Following QC, nuclei samples had average gene and feature counts per cell of 2849 and 1496 respectively, while those counts for cells were 4893 and 1966, respectively. The final integrated dataset consisted of 220,752 nuclei and 49,723 cells representative of 15 major cell types (**Fig. 1C**). Cell identities were validated by expression of cell specific marker genes (**Fig. 1D, Fig. S3**) and transcriptional signatures (**Fig. S4-6**). Cell types identified in both snRNAseq and scRNAseq datasets included fibroblasts, endothelial cells, myeloid cells, pericytes, smooth muscle cells, T- and NK-cells, neurons/glia, and B-cells. A notable benefit of snRNAseq is the ability to obtain reads from additional cell types that are not efficiently recovered from enzymatically digested tissue including cardiomyocytes, adipocytes, endocardial cells, lymphatics, epicardial cells, and mast cells (**Fig. 1E**). Quantification of major cell type distribution revealed that the percentages of fibroblasts, myeloid cells, and NK- and T-cells were increased in DCM compared to non-diseased tissues (**Fig. S7**).

The analyzed dataset was powered to investigate the influence of age, sex, and disease state on gene expression. Differential expression analysis using pseudobulk and single cell approaches demonstrated that disease state had the most powerful influence on differential gene expression across cell types (**Fig. 2A**). Substantially fewer differentially expressed genes were detected comparing non-diseased males and females, the majority of which were located on the X and Y chromosomes including XIST, TSIX, and TTTY genes (**Fig. 2B**). Moreover, comparison of non-diseased young and old subjects did not reveal clear differences (**Fig. 2C**). With respect to disease state, cardiomyocytes, myeloid cells, fibroblasts, endothelial cells, and endocardial cells displayed the greatest differences in gene expression in both snRNAseq and scRNAseq datasets (**Fig. 2A**). Pseudobulk and Seurat differential expression also demonstrate substantial overlap in the genes found to be differentially expressed, with pseudobulk identifying a larger number of differentially expressed genes (**Fig. S8**). As such, we focused our downstream analysis on these cell types.

**Figure 2.**
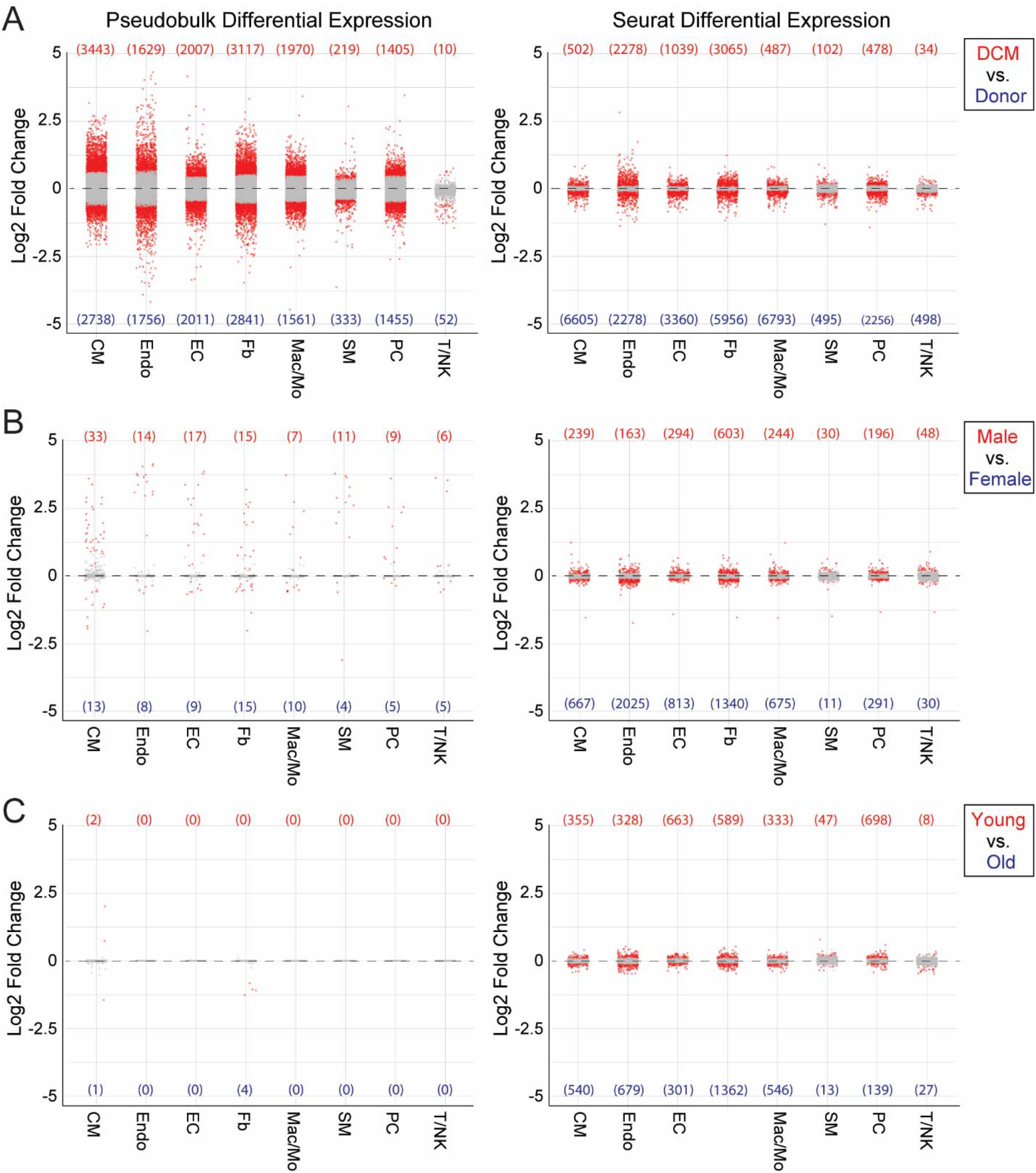
Differential influence of disease state, sex, and age on cell type specific gene expression. Dot plots showing pseudobulk (DESeq2, left) and single cell (Seurat, right) based differential gene expression across major cell populations. Differential expression was calculated from single nucleus RNA sequencing data for disease (**A**, Donor vs DCM), sex (**B**, male vs female), and age (**C**, young vs old) is shown. Differential expression analysis for age was calculated by dividing samples into younger (<52 years old) and older (>52 years old) groups based on median age. Genes with adjusted p-value <0.05 colored in red and genes with adjusted p-value >0.05 are colored in grey. Number of upregulated and downregulated genes with adjusted p-value <0.05 per cell type is displayed in parenthesis.

### Cardiomyocytes phenotypically converge in dilated cardiomyopathy

Principal component analysis of pseudobulk data indicated that disease state and sex influenced gene expression profiles of cardiomyocytes. Overlaying age distribution onto the PCA plot did not suggest a strong relationship with age across all cardiomyocytes although differential expression analysis identified a small number of genes differentially expressed between older vs. younger individuals (**Fig. 3A, Fig. S9**). Pseudobulk differential expression analysis between male and female subjects indicated robust differences in genes encoded on the X and Y chromosome, possibly accounting for separation observed by PCA (**Fig. S9**). Differential expression analysis by pseudobulk and single cell approaches across disease state revealed a large number of genes significantly upregulated (NPPA, RYR1, EGR2) and downregulated (EDNRB, MYH6, CKM) in DCM samples compared to non-diseased donors (**Fig. 3B, Fig. S9**). Pathway analysis identified multiple differentially regulated pathways including VEGFA signaling, metabolism, and proteasome degradation. Comparison of enriched pathways between clusters identified distinct pathways associated with individual cell states. (**Fig. S10**). Transcription factor enrichment analysis was also performed on select states of cardiomyocytes associated with DCM. The ADGRL3 cell state was found to have enrichment for targets of specific transcription factors including STAT3, CEBPD, and SMARCD1, while targets of ZNF217, WT1, and AR, among others were identified in analysis of the NPPA/NPPB cell state (**Fig. S10**).

**Figure 3.**
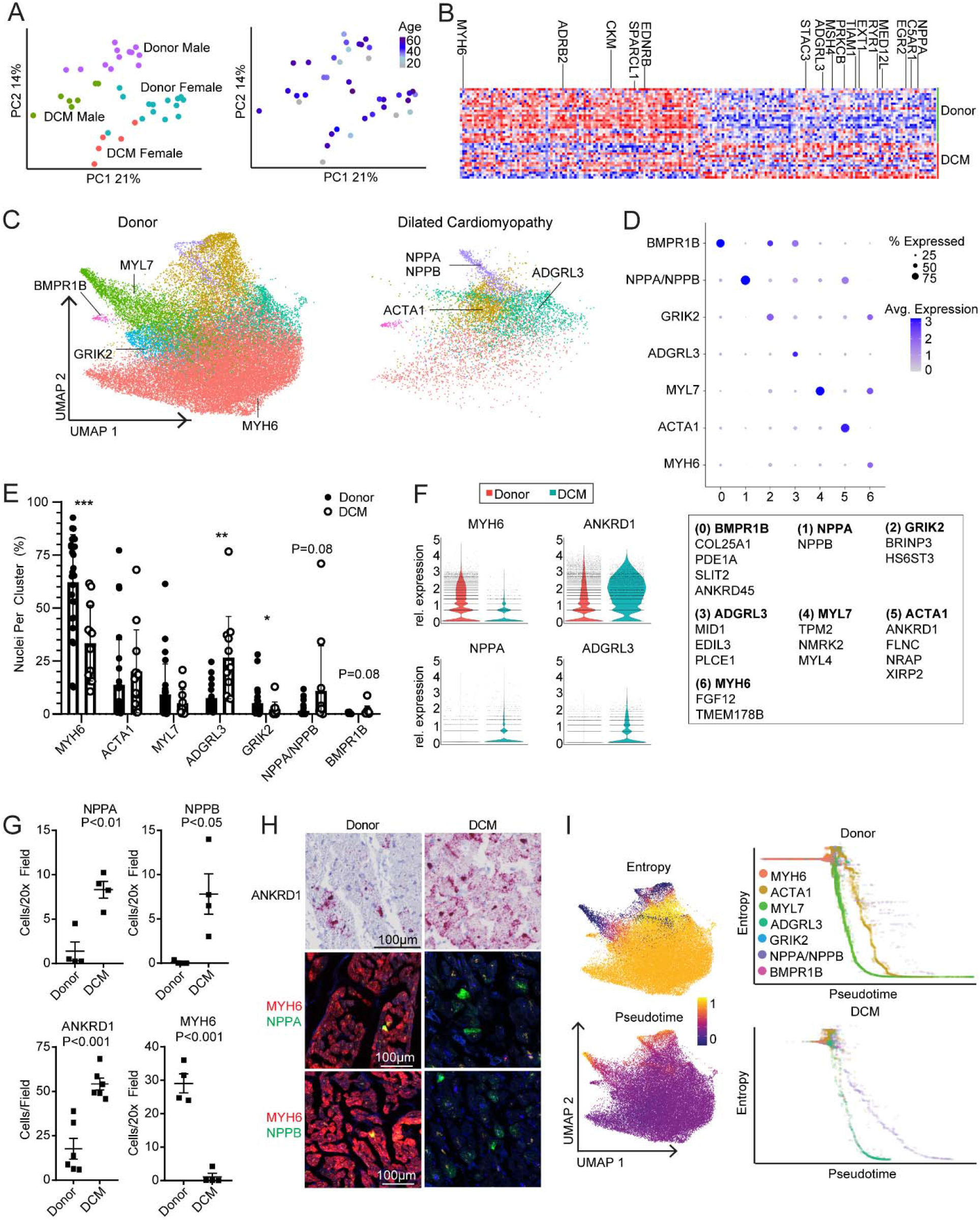
Acquisition of disease associated cardiomyocyte states in dilated cardiomyopathy. **A,** Principal component analysis (PCA, DESeq2) plots of cardiomyocyte pseudobulk single nucleus RNA sequencing data colored by sex and disease state (left) and age (right). Each data point represents an individual subject. **B,** Heatmap displaying the top 100 upregulated and downregulated genes ranked by log2 fold-change comparing donor control to dilated cardiomyopathy (DCM). Differentially expressed genes were derived from the intersection of pseudobulk (DESeq2) and single cell (Seurat) analyses. **C,** Unsupervised clustering of donor and DCM cardiomyocytes within the integrated dataset split by disease state. Major cardiomyocyte states are labeled. **D,** Dot plot displaying the z-scores for transcriptional signatures that distinguish cardiomyocyte states (genes selected by enrichment in Seurat differential expression analysis, listed in box below plot). **E,** Distribution of cardiomyocyte states by cluster (*<0.05, **<0.01, ***<0.001, Welch’s T-test, two-tailed, lines represent mean and standard deviation). **F,** Violin plots of MYH6, ANKRD1, NPPA, and ADGRL3 expression in donor control and DCM cardiomyocytes. **G,** Quantification of the number of cardiomyocytes expressing ANKRD1, MYH6, NPPA, and NPPB mRNA in donor control and DCM samples (p-value from Welch’s T-Test, two-tailed, lines represent mean and standard deviation). **H,** Representative RNA *in situ* hybridization images (RNAScope) of indicated genes. **I.** Palantir pseudotime trajectory analysis of cardiomyocytes showing entropy and pseudotime scores overlaid on the UMAP projection (left). Entropy vs pseudotime plots of donor and DCM cardiomyocytes identifying differing trajectories of healthy and disease associated cardiomyocyte states (right).

Unsupervised clustering identified 7 cardiomyocyte states with differing gene expression signatures **(Fig. 3C-D, Fig. S10)**. Cardiomyocytes from donor samples existed in all 7 states marked by MYH6, MYL7, GRIK2, NPPA/NPPB, ADGRL3, and ACTA1 expression. Intriguingly, cardiomyocytes from DCM samples displayed a striking bias towards clusters marked by NPPA/NPPB and ADGRL3 expression. There was a marked reduction in the MYH6 cardiomyocyte clusters (**Fig. 3C,E**). In addition to changes in cell distribution, we also observed a decrease in MYH6 expression and increases in ANKRD1, NPPA, and ADGRL3 expression in DCM (**Fig. 3F**). To validate shifts in cardiomyocyte state and gene expression in DCM at the tissue level, we performed RNA *in situ* hybridization. Compared to donor controls, we observed significant increases in NPPA, NPPB, and ANKRD1 expressing cells and significant reduction in MYH6 expressing cells in DCM (**Fig. 3G-H**).

To explore the temporal relationship between cardiomyocyte states, we performed pseudotime trajectory analysis using Palantir.^11^ We calculated pseudotime and entropy values for each cardiomyocyte cluster to predict putative states of cell differentiation (**Fig. 3I, Fig. S10**). We plotted entropy versus pseudotime values for each cell and superimposed cluster designations. Donor cardiomyocytes were predicted to contain two highly differentiated cell states marked by MYL7 and ACTA1 expression. In contrast, DCM samples displayed two distinct highly differentiated cell states marked by ARGRL3 and NPPA/NPPB expression (**Fig. 3I**). Collectively, these observations suggest a convergence towards disease associated cardiomyocyte phenotypes in DCM.

### Expansion of monocytes and shift in macrophages toward diverse inflammatory states

Macrophages, monocytes, and dendritic cells are increasingly studied in mouse models of cardiac injury and heart failure.^12–16^ We identified large populations of macrophages, monocytes, and dendritic cells in donors and DCM subjects (**Fig. 1C,E**). Principal component analysis of pseudobulk data indicated that disease state and sex had the greatest effect differential gene expression. We did not identify an obvious relationship with age (**Fig. 4A, Fig. S9**). Differential expression analysis by pseudobulk and single cell approaches across disease state revealed a large number of genes significantly upregulated (CCL3, CD1C, NLRP3) and downregulated (VSIG4, FCGBP, MCM2) in DCM samples compared to non-diseased donors (**Fig. 4B**). Similar to cardiomyocytes, pseudobulk differential expression analysis between male and female subjects indicated robust differences in a small number of genes encoded on the X and Y chromosomes, including XIST, JPX, and TTTY10 (**Fig. S9).** Pathway analysis identified upregulation of multiple pathways in DCM samples including IL-18, PI3K-AKT-mTOR, IL-3, and MAPK signaling, while cell cycle and metabolism pathways were found to be downregulated in DCM **(Fig. S11).**

**Figure 4.**
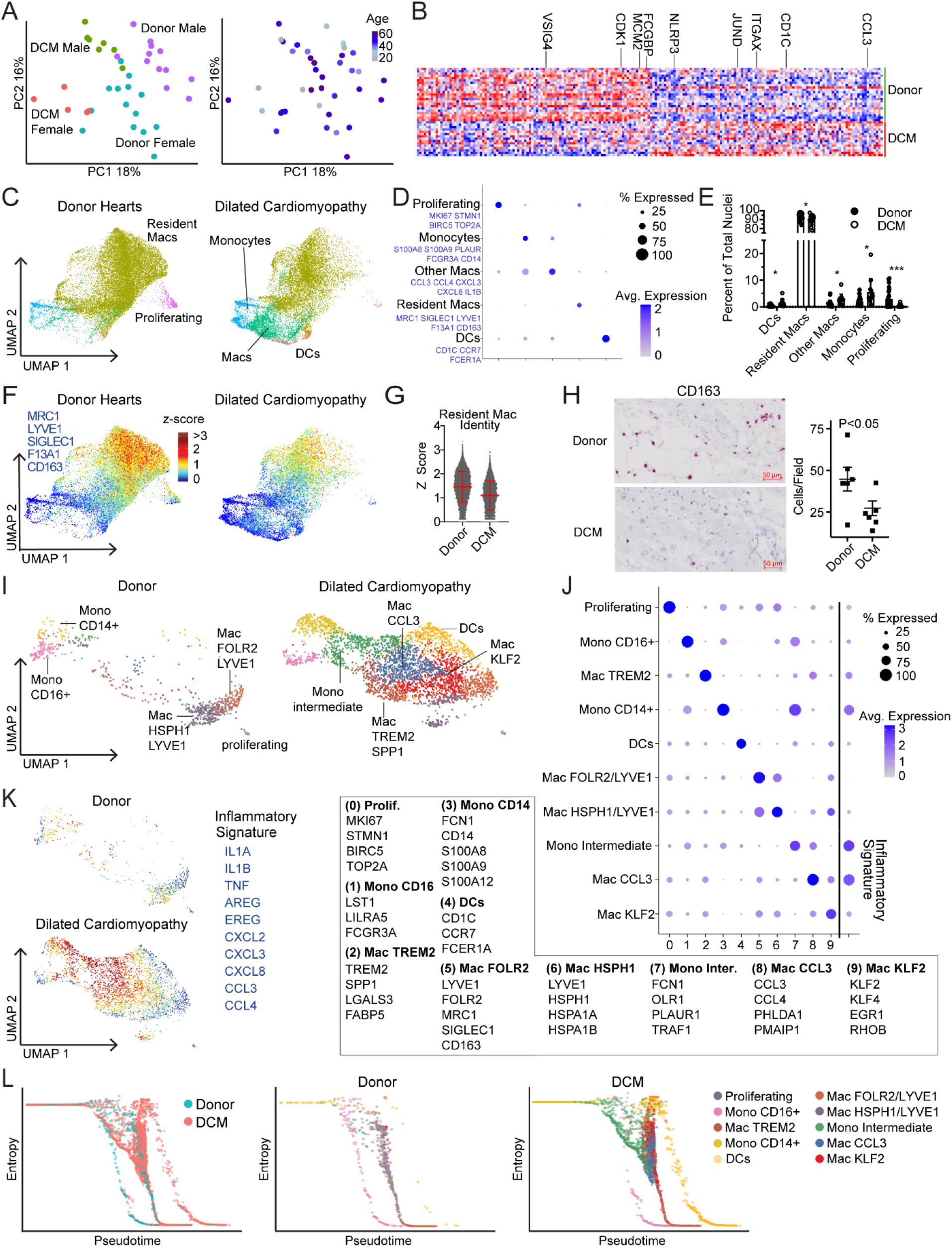
Dilated cardiomyopathy is associated with shifts in macrophage composition and gene expression favoring inflammatory populations. **A,** Principal component analysis (PCA, DESeq2) plots of monocyte, macrophage, and dendritic cell pseudobulk single nucleus RNA sequencing data colored by sex and disease state (left) and age (right). Each data point represents an individual subject. **B,** Heatmap displaying the top 100 upregulated and downregulated genes ranked by log2 fold-change comparing donor control to dilated cardiomyopathy (DCM). Differentially expressed genes were derived from the intersection of pseudobulk (DESeq2) and single cell (Seurat) analyses. **C,** Unsupervised clustering of monocytes, macrophages, and dendritic cells within the integrated dataset split by disease state. Major cell states are labeled. **D,** Dot plot displaying the z-scores for transcriptional signatures that distinguish monocyte, macrophage, and dendritic cell populations (genes selected by enrichment in Seurat differential expression analysis are listed in blue). **E,** Distribution of myeloid states by cluster (*<0.05, ***<0.001, Welch’s T-test, two-tailed, lines represent mean and standard deviation, derived from single nucleus data). **F-G,** Z-score feature plot (F) and violin plot (G) of the tissue resident macrophage signature split by disease state. **H,** Representative RNA *in situ* hybridization images (RNAScope) for CD163 (red, blue: hematoxylin) and quantification of CD163+ cells in donor and DCM samples (p-value from Welch’s T-Test, two-tailed, lines represent mean and standard deviation). CD163 is a marker of tissue resident macrophages. **I,** UMAP projection of unsupervised re-clustering of myeloid cells from the single cell RNA sequencing dataset. Major cell states are labeled. **J,** Dot plot displaying the z-scores for transcriptional signatures that distinguish each monocyte, macrophage, and dendritic cell state (genes selected by enrichment in Seurat differential expression analysis, listed in box below plot). **K,** Z-score feature plot overlaying an inflammatory gene expression signature (genes in blue) on the single cell RNA sequencing UMAP projection split by disease state. **L,** Palantir pseudotime trajectory analysis of myeloid single cell RNA sequencing data. Entropy vs pseudotime plots split by disease state identify major cell trajectories (non-classical monocytes, resident macrophages, dendritic cells). Inflammatory cell states that emerge in DCM have high entropy and low pseudotime values suggesting an intermediate state of differentiation.

Unsupervised clustering of the integrated reference dataset revealed the presence of large numbers of macrophages, and smaller populations of monocytes, dendritic, and proliferating cells. We identified two populations of macrophages including a subset that expressed tissue resident markers (MRC1, SIGLEC1, CD163, LYVE1, F13A1).^17–19^ Compared to donor controls, we observed a reduction in resident and proliferating macrophages and expansion of monocytes, dendritic cells, and other macrophages in DCM subjects (**Fig. 4C-E, Fig. S11**). We also observed a reduction in the tissue resident macrophage signature in DCM (**Fig. 4F-G**). RNA *in situ* hybridization confirmed reduction in CD163+ cells DCM samples and compared to donor controls (**Fig. 4H**).

Visualization of snRNA-seq and scRNA-seq data within the integrated object indicated a bias in recovered cell populations. While each dataset contained all of the identified cell types, the scRNAseq dataset displayed a bias towards monocytes, dendritic cells, and non-resident macrophages. The snRNAseq dataset contained a substantially larger number of resident macrophages (**Fig. S11**). To evaluate the diversity of monocytes, dendritic cells, and non-resident macrophages, we chose to focus on the scRNAseq data. Unsupervised clustering revealed the presence of discrete monocyte (CD14+, CD16+, intermediate), macrophages (CCL3, TREM2, KLF2, LYVE1) and dendritic cell populations (**Fig. 4I-J).** We observed shifts in monocyte, macrophage, and dendritic cell composition between donor and DCM groups. Donor samples contained classical (CD14+) and non-classical (CD16+) monocytes as well as 2 populations of LYVE1+ resident macrophages that differed based on the expression of FOLR2, MRC1, SIGLEC1, and HSPH1. Compared to donors, DCM samples displayed reduced numbers of resident macrophages and a greater number of intermediate monocytes, dendritic cells, and three additional macrophage populations characterized by the expression of CCL3, TREM2, and KLF2 (**Fig. S11**). Classical and intermediate monocytes and macrophages marked by CCL3, TREM2, and KLF2 expressed robust levels of inflammatory mediators including IL1A, IL1B, TNF, AREG, EREG, and multiple chemokines (**Fig. 4J-K**). Comparison of enriched pathways across cell states identified enrichment of unique pathways in individual states including enrichment of inflammatory IL10 and IFNγ signaling in the CCL3 state, and enrichment of IFN α,β, and γ in the intermediate monocyte state. Transcription factor analysis identified enrichment of targets of transcription factors including CLOCK, RELA, MYB, and IRF8 in the inflammatory macrophage states (CCL3, KLF2, and TREM2) (**Fig. S12**).

To infer the differentiation state of monocyte, dendritic cell and macrophage populations, we utilized Palantir. Calculation of pseudotime and entropy values demonstrated that CD14+ monocytes represented the most progenitor-like state. CD16+ monocytes, dendritic cells, and resident macrophages represented the most differentiated cells, each with distinct trajectories. Compared to donors, we observed an accumulation of cells with intermediate differentiation states along the macrophage trajectory in DCM samples. Superimposing cluster identities revealed that these cells belonged to the intermediate monocyte, TREM2, CCL3, and KLF2 clusters suggesting that they are monocyte-derived (**Fig. 4L**). These data provide a link between CD14+ monocytes, monocyte-derived macrophage diversity and inflammation in DCM.

### Fibroblasts diversify in dilated cardiomyopathy

We identified a large population of cardiac fibroblasts that displayed a dramatic shift in gene expression in DCM as compared to donor controls (**Fig. 1C)**. Principal component analysis demonstrated that variability across samples was driven by disease state and sex with less influence of age. (**Fig. 5A, Fig. S9**). Differential expression analysis by pseudobulk identified subtle differences in gene expression by age (**Fig. S9**). Pseudobulk and single cell differential expression analysis identified a large number of genes significantly upregulated (LEF1, NFATC2, PTCH2) and downregulated (GPX3, FGF10, FGF7) in DCM samples compared to non-diseased donors. Pathway analysis identified upregulation (MAPK, EGF, WNT, BDNF signaling) and downregulation (metabolism, biosynthesis) of multiple pathways in DCM (**Fig. S13**). Similar to cardiomyocytes and macrophages, pseudobulk differential expression analysis between male and female subjects indicated robust differences in a small number of genes encoded on the X and Y chromosomes including XIST, JPX, and ZFYAS1 (**Fig. S9**).

**Figure 5.**
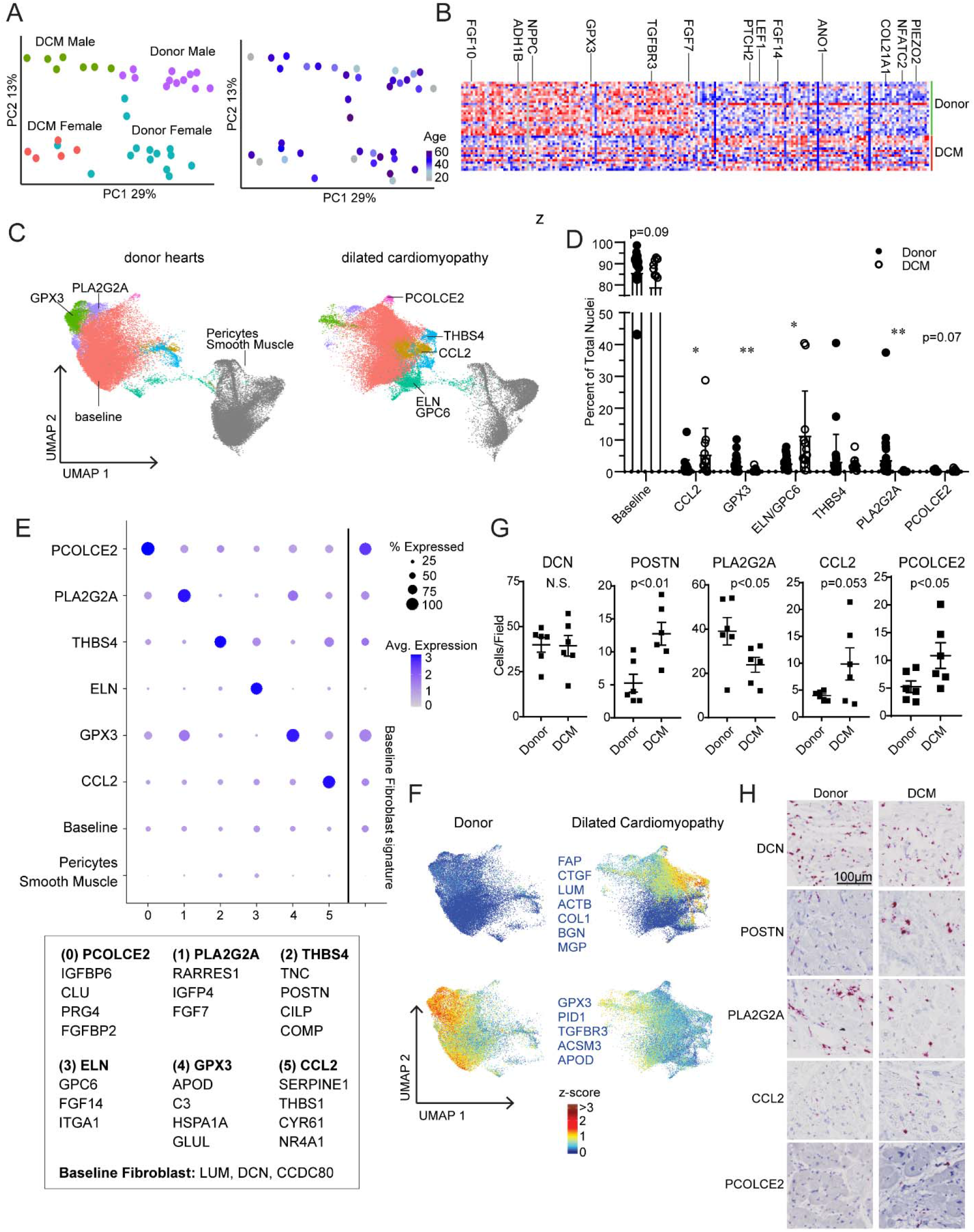
Phenotypic shifts and emergence of disease associated fibroblasts in dilated cardiomyopathy. **A,** Principal component analysis (PCA, DESeq2) plots of fibroblast pseudobulk single nucleus RNA sequencing data colored by sex and disease state (left) and age (right). Each data point represents an individual subject. **B,** Heatmap displaying the top 100 upregulated and downregulated genes ranked by log2 fold-change comparing donor control to dilated cardiomyopathy (DCM). Differentially expressed genes were derived from the intersection of pseudobulk (DESeq2) and single cell (Seurat) analyses. **C,** Unsupervised clustering of donor and DCM fibroblasts within the integrated dataset split by disease state. Major fibroblast states are labeled. Pericytes and smooth muscle cells are displayed in grey. **D,** Distribution of fibroblast states by cluster (*<0.05, **<0.01, Welch’s T-test, two-tailed, lines represent mean and standard deviation). **E,** Dot plot displaying the z-scores for transcriptional signatures that distinguish fibroblast states (genes selected by enrichment in Seurat differential expression analysis, listed in box below plot). **F,** Z-score feature plot of transcriptional signatures associated with DCM (top) and with donor (bottom) fibroblast states. Plot is split by disease state. DCM fibroblasts are enriched in genes associated with activation. Enriched genes (blue) were defined using Seurat differential gene expression analysis. **G,** Quantification of the number of cell expressing DCN, POSTN, PLA2G2A, CCL2, and PCOLCE2 mRNA in donor control and DCM samples (p-value from Welch’s T-Test, two-tailed, lines represent mean and standard deviation). **H,** Representative RNA *in situ* hybridization images (RNAScope) of indicated genes (red). Blue hematoxylin.

Unsupervised clustering of the integrated reference dataset revealed multiple distinct populations of fibroblasts (**Fig. 5C**). The majority of fibroblasts in both donor and DCM hearts displayed a conserved gene expression signature characteristic of fibroblasts. We identified two fibroblast subpopulations primarily present in donor controls that expressed PLA2G2A and GPX3, respectively. In the context of DCM, we observed expansion of additional fibroblast subpopulations characterized by the expression of PCOLCE2, THBS4, CCL2, and ELN/GPC6. Interestingly, the THBS4 cluster contained two signatures marked by POSTN and TNC expression with POSTN selectively found in DCM samples (**Fig. 5C-E, Fig. S13**). Fibroblasts in DCM hearts displayed a robust activation signature that included FAP, CTGF, LUM, ACTB, COL1A1, BGN, and MGP expression. Donor fibroblasts selectively expressed GPX3, PID1, TGFBR3, ACSM3, and APOD (**Fig. 5F**). Palantir identified fibroblasts marked by THBS4, CCL2, and ELN expression as the most differentiated cell states based on low entropy and high pseudotime values. All other fibroblasts appeared to exist in a state of high entropy suggesting significant plasticity within these populations (**Fig. S13**). Additional pathway analysis comparing populations identified distinct enrichment of pathways in individual cell states, including pathways involved in protein translation and transport in the CCL2, GPX3, and THBS4 populations and enrichment of striated muscle contraction associated pathways in the unactivated population (**Fig. S13**). Transcription factor analysis on select populations associated with DCM identified enrichment of targets of specific transcription factors including CREM, NELFA, EGR1, and TCF7 in the CCL2 population and TRIM28, RUNX1, BRD4, CJUN, and MYC in the THBS4 population (**Fig. S14**).

We validated shifts in fibroblast composition between donor controls and DCM hearts using RNA *in situ* hybridization. The overall numbers of fibroblasts (marked by DCN expression) remained similar between donor control and DCM hearts. Interesting, we observed that fibroblast subpopulations were located either within the interstitial space between cardiomyocytes (PCOLCE2, CCL2, POSTN), adjacent to distal vasculature (PLA2G2A), or surrounding epicardial coronary arteries (ELN). The number of POSTN fibroblasts was significantly increased in DCM samples. We also identified strong trends for reduced PLA2G2A fibroblasts (p=0.06) and increased CCL2 fibroblasts (p=0.08) in DCM. (**Fig. 5G-H, Fig. S13**).

Unsupervised clustering of the integrated reference dataset subtle changes in pericytes and smooth muscle cells. Principal component analysis demonstrated that variability across samples was driven by disease state and sex with little influence of age. Pseudobulk and single cell differential expression and pathway analyses identified genes and pathways enriched in DCM pericytes and smooth muscles cells compared to non-diseased donors. We did not observe distinct subpopulations of pericytes or smooth muscles in any examined condition (**Fig. S15**).

### Endothelial cell populations display shifts in global gene expression

Endothelial cells within the heart include arterial, venous, capillary, lymphatic, and endocardial cells. Principal component analysis of artery, vein, and capillary pseudobulk data identified disease state and sex as distinguishing features with less impact from age (**Fig. 6A, Fig. S9**). Differences between male and female subjects were driven by a small number of genes encoded on the X and Y chromosomes (**Fig. S9**). Pseudobulk bulk and single cell differential expression analysis in vascular endothelial cells identified a large number of genes significantly upregulated (EGR1, PLXNA4, NOX4, PDE4) and downregulated (SIPR3, TBX3, FABP5) in DCM samples compared to non-diseased donors (**Fig. 6B**).

**Figure 6.**
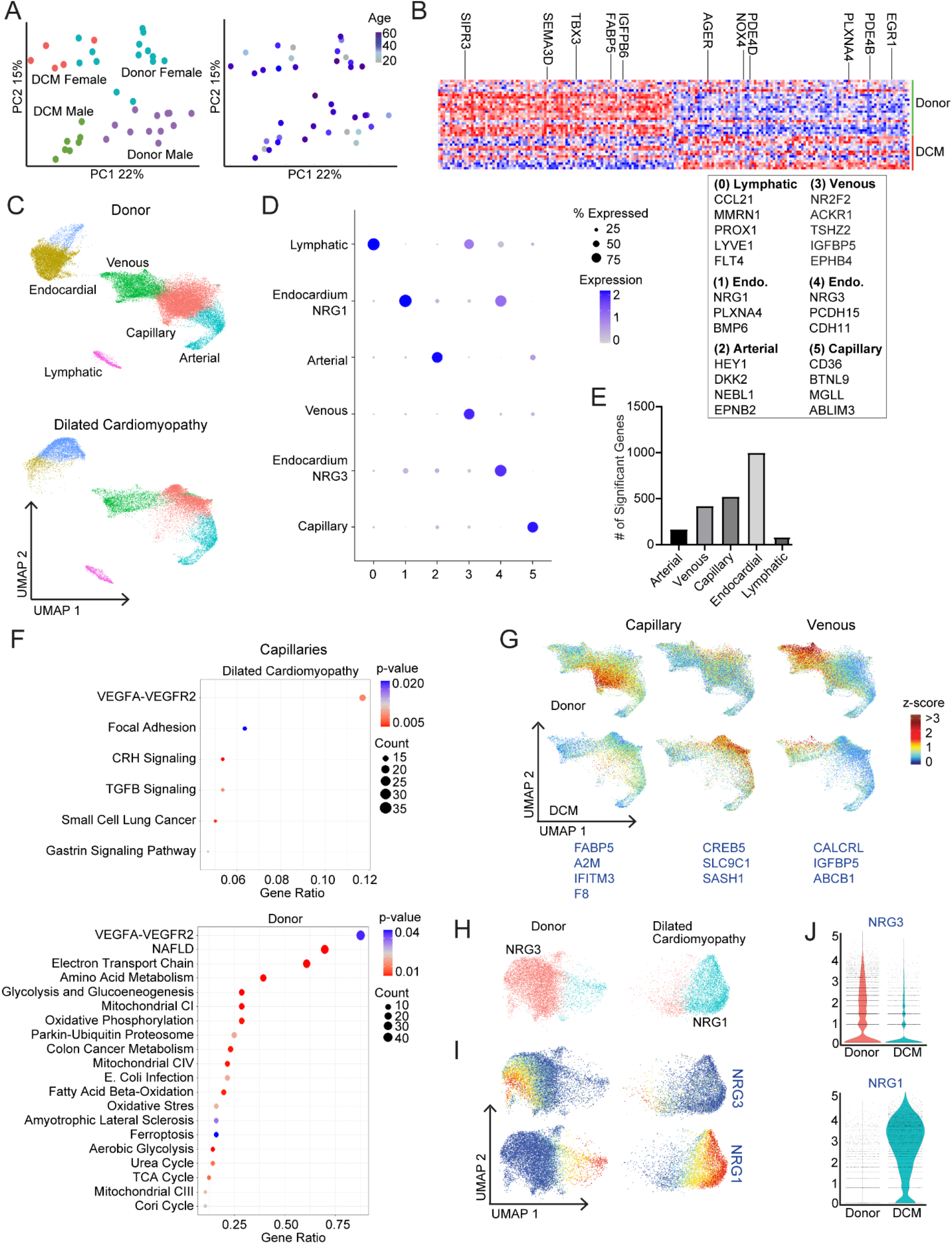
Endothelial and endocardial cells exhibit global gene expression shifts in dilated cardiomyopathy. **A,** Principal component analysis (PCA, DESeq2) plots of vascular endothelial cell pseudobulk single nucleus RNA sequencing data colored by sex and disease state (left) and age (right). Each data point represents an individual subject. **B,** Heatmap displaying the top 100 upregulated and downregulated genes ranked by log2 fold-change comparing donor control to dilated cardiomyopathy (DCM). Differentially expressed genes were derived from the intersection of pseudobulk (DESeq2) and single cell (Seurat) analyses. **C,** Unsupervised clustering of donor and DCM vascular endothelial cells within the single nucleus RNA sequencing dataset split by disease state. Major endothelial states are labeled. **D,** Dot plot displaying z-scores for transcriptional signatures that distinguish endothelial cell populations (genes selected by enrichment in Seurat differential expression analysis, genes listed in the box to right of plot). **E,** Bar graph of the number of differentially expressed genes per endothelial population (intersection of DESeq2 and Seurat differential expression analyses, adjusted p<0.05, log2FC>0.1). **F,** WikiPathways analysis identifying pathways enriched in donor and DCM capillary endothelial cells. **G,** Z-score feature plots of transcriptional signatures associated with donor and DCM groups in capillary and venous endothelial cells split by disease state. Genes (blue) were selected by enrichment in the differential expression analyses. **H,** Unsupervised clustering of donor and DCM endocardial cells (single nucleus RNA sequencing data) split by disease state. **I.** UMAP feature plots of NRG1 and NRG3 split by disease state. **J.** Violin plots displaying NRG1 and NRG3 expression in endocardial cells from donor and DCM samples.

Within the integrated reference object, the snRNAseq dataset contained all major endothelial cell populations whereas the scRNAseq dataset displayed a bias towards arterial, venous, and capillary endothelial cells. Few endocardial or lymphatic cells were recovered from scRNAseq data. Utilizing RNA in situ hybridization, we visualized expression of arterial (CLIC3), venous (ACKR1), capillary (BTNL9), and lymphatic (CCL21) markers identified from Seurat differential expression analysis (**Fig. S16**). To examine whether DCM is associated with changes in endothelial cell diversity, we performed unsupervised clustering of the snRNAseq and scRNAseq datasets. This analysis did not reveal further diversification of arterial, venous, capillary, lymphatic, or endocardial cells. Instead, we observed global shifts in gene expression across both datasets (**Fig. 6C-D, Fig. S16**).

Pseudobulk and single cell differential gene expression analysis of snRNAseq data demonstrated that capillaries, veins, and endocardial cells displayed the greatest number of differentially expressed genes comparing donor control and DCM conditions (**Fig. 6E**). Vascular endothelial cells and endocardial cells displayed distinct transcriptional signatures and pathways enriched in DCM samples. Capillaries and venous endothelial cells exhibited distinct signatures in DCM samples as compared to non-diseased donors. Upregulation of CREB5, SLC9C1, and SASH1 was observed in capillaries of DCM samples while FOS and DUSP1 was upregulated in both venous and capillaries in DCM samples. Donor capillaries exhibited upregulation of FABP5 and IFITM3 and donor venous cells exhibited strong enrichment of CALCRL and IGFBP5. Pathway analysis identified downregulation of metabolic pathways in both venous and capillary endothelial cells in DCM samples. We also identified enrichment of TGF-beta, CRH, and Kit signaling and the hallmark pathway of cardiovascular disease in venous endothelial cells in DCM. In capillaries, we identified enrichment of VEGFA, CRH, TGF-beta, and gastrin signaling in DCM as compared to donors (**Fig 6F-G, Fig. S16**).

Within endocardial cells, we observed a large number of genes to be significantly upregulated (BMP4, NRG1, NRP2) and downregulated (IGFBP6, CD55, ITGA6) in DCM samples compared to non-diseased donors (**Fig. S17**). Endocardial cells displayed a striking shift in gene expression between donor controls and DCM resulting in independent clustering of endocardial cells across disease state (**Fig. 6H**). Donor endocardial cells expressed NRG3. Endocardial cells from DCM samples display strong upregulation of NRG1 and reduced NRG3 expression (**Fig. 6I-J**). Pathway analysis identified enrichment of PI3K-AKT, MAPK, EGF, hedgehog, and TGF-beta signaling as well as hallmarks of heart development in DCM samples while VEGFA signaling, and metabolic pathways were enriched in donor endocardial cells (**Fig. S17**).

Additional pathway analysis comparing NRG3 to NRG1 populations identified enrichment of muscle contraction pathways in the NRG3 population that are absent in the NRG1 population as well as enrichment of FGFR1 signaling in the NRG1 population. Transcription factor analysis identified enrichment of targets of transcription factors including FOXA2, STAT3, TCF4, GATA2, and FOXM1 in the NRG1 population as well as TBX20, RELA, SOX9, and SCL/TAL1 in the NRG3 population (**Fig. S17**).

## Discussion

Single cell technologies offer powerful new tools to dissect cell types that reside within healthy and diseased tissues. Recently, these approaches were leveraged to provide a deeper understanding of the cellular composition of the healthy human heart.^6,7^ While considerable interest exists, only limited data is available to decipher how the cellular and transcriptional landscape of the heart is impacted by disease.^20,21^

This study represents the largest-scale single cell analysis of the healthy and failing human heart to date. Using an approach that integrated snRNAseq and scRNAseq data from 45 individuals encompassing 220,752 nuclei and 49,723 cells, we identified 15 major cardiac cell types, uncovered cell type specific transcriptional programs, and revealed the emergence of cell states associated with heart failure. We did not detect significant differences in cellular composition related to age or sex. However, this does not exclude the possibility that aging and/or sex may have impacts that were not readily identified by single cell analysis. We observed considerable variation in how different cardiac cell populations responded to heart failure. Cardiomyocytes converged towards a common disease associated cell state, while fibroblasts and myeloid cells underwent dramatic diversification including the acquisition of disease specific phenotypes. In contrast, endothelial cells, endocardial cells and pericytes displayed global transcriptional shifts without changes in cell complexity.

Previous studies examining differences across cardiac chambers have identified evidence of cardiomyocyte heterogeneity in the healthy heart.^6,7^ We identified multiple transcriptionally distinct cardiomyocyte states within the LV of non-diseased donors and DCM patients. Donor hearts contained 7 cardiomyocyte states marked by MYH6, MYL7, GRIK2, NPPA/NPPB, ADGRL3, and ACTA1 expression. DCM cardiomyocytes uniformly expressed high levels of ARKD1, contained few MYH6 or GRIK2 expressing cardiomyocytes, and instead, were enriched for states identified by NPPA/NPPB and ADGRL3 expression. Interesting, ANKRD1 expression was recently found to be enriched in cardiomyocytes from patients with adolescent versus pediatric DCM.^20^ Pseudotime trajectory analysis identified three highly differentiated cardiomyocyte states (MYL7, ACTA1, NPPA/NPPB) in donor hearts. Surprisingly, we observed a two highly differentiated cardiomyocyte state in DCM marked by ADGRL3 and NPPA/NPPB expression. These observations suggest that cardiomyocytes converge towards a common disease associated states in DCM. Further understanding of the instructive cues and parental cardiomyocyte populations that give rise to ADGRL3 and NPPA/NPPB expressing cardiomyocytes may provide new insights and opportunities to intervene in the pathogenesis of human heart failure.

We observed striking transcriptionally changes in non-cardiomyocyte populations (fibroblasts, macrophages, endothelial cells, endocardial cells) between healthy controls and DCM samples. Prior snRNAseq studies have reported astounding diversity amongst fibroblasts in the healthy human heart.^6,7,22,23^ Fibroblasts are known to expand in heart failure and acquire an activated phenotype characterized by the expression of fibroblast activated protein (FAP) and periostin (POSTN).^24–30^ While previous single cell studies have identified cardiac fibroblast subsets in the healthy human heart, little is known regarding how these populations are influenced by disease. We identify multiple distinct fibroblast populations in both healthy and diseased samples with differing transcriptional signatures and spatial distribution including elastin (ELN) expressing macrophages located within the media of coronary arteries. Fibroblasts marked by POSTN, CCL2, and PCOLCE2 were enriched in DCM, while GPX3 and PLA2G2A macrophages were enriched in donor controls. In addition, we identified an activation signature that included FAP, CTGF, LUM, ACTB, COL1A1, BGN, and MGP that was selectively expressed in fibroblasts from DCM hearts. These findings provide further evidence that phenotypic shifts in fibroblasts are a hallmark of heart failure.

Heterogeneity of myeloid populations including macrophages is increasingly appreciated to contribute to the variety of cardiac pathologies including heart failure.^31–36^ The majority of these studies have focused on mouse models with only targeted validation in human specimens.^14,15,37,38^ Consistent with small animal models, we observe a variety of monocyte, macrophage, and dendritic cell populations within the human heart. The abundance of macrophages expressing a tissue resident signature is reduced in DCM. We also observed an emergence of monocyte and macrophage populations expressing inflammatory mediators in the failing heart. Cell trajectory analysis predicted that many of these inflammatory populations represented intermediate states derived from CD14+ monocytes. The number of proliferating macrophages was reduced in DCM, consistent with the concept that self-replication maybe a trait of tissue resident macrophages.

While scRNAseq and snRNAseq provided sufficient resolution to identify major perivascular populations (arteries, veins, capillaries, pericytes, smooth muscle cells, lymphatics, endocardial cells), we did not observe additional diversity within these populations. However, we did uncover global shifts in gene expression within each of these populations between control and DCM specimens. Previous studies have identified similar shifts in global endothelial cell expression but were unable to parse contributions from each major endothelial cell type. Endocardial cells displayed robust numbers of differentially expressed genes between control and DCM specimens. NRG1 and NRG3 were exclusively expressed in DCM and control endocardial cells, respectively. Interestingly, mouse studies identified that cardiomyocyte specific loss of NRG3 receptors (ErbB2, ErbB4) results in spontaneous heart failure suggesting a potential role for NRG3 in regulating cardiac homeostasis.

snRNAseq captured cell types that are difficult to recover from enzymatically digested tissue including cardiomyocytes, adipocytes, mast cells, epicardium, endocardium, and lymphatics. Using data integration and reference mapping, we were able to effectively combine snRNAseq and scRNAseq data and identify at least 15 major cardiac cell populations. Current scRNAseq datasets exploring human heart failure are small and lack the sample size necessary to elucidate the impact of disease on common and rare cardiac cell types.^20,21^ scRNAseq data provided greater depth at the expense of biased cell recovery. For example, within myeloid cells, scRNAseq data was biased towards monocytes and intermediate macrophage populations with fewer resident macrophages recovered. These datasets were leveraged to provide additional granularity into monocytes and inflammatory macrophage populations.

This study is not without limitations. We categorized DCM patients into a single cohort based on the lack of underlying coronary artery disease. It is likely that the exact etiology of DCM contributes to shifts in cell diversity and transcriptional state. Our data set only includes transcriptomic information. Addition of cell surface protein expression and chromatin accessibility information may offer additional resolution. In conclusion, this study represents the largest analysis of the cellular and transcriptomic landscape of the healthy and failing human heart to date. We provide valuable insights into how cardiac cell populations change during heart failure including the emergence of disease specific cell states. These data provide a valuable resource that will open up new areas of investigation and opportunities for therapeutic development and innovation.

## Acknowledgements

This project was made possible by funding provided from the NHLBI (R01 HL138466, R01 HL139714, R01 HL151078, T32 HL134635, T32 HL125241), Leducq Foundation Network (#20CVD02), Burroughs Welcome Fund (1014782), Children’s Discovery Institute of Washington University and St. Louis Children’s Hospital (CH-II-2015-462, CH-II-2017-628, PM-LI-2019-829), Foundation of Barnes-Jewish Hospital (8038-88), as well as Aging Biology Foundation awarda to M.N.A. Histology was performed in the DDRCC advanced imaging and tissue analysis core supported by Grant #P30 DK52574. Imaging was performed in the Washington University Center for Cellular Imaging (WUCCI) which is funded, in part by the Children’s Discovery Institute of Washington University and St. Louis Children’s Hospital (CDI-CORE-2015-505 and CDI-CORE-2019-813) and the Foundation for Barnes-Jewish Hospital (3770). We acknowledge the McDonnel Genome Institute for their assistance in designing and performing the micro array and RNA sequencing analysis as well as the Mallinckrodt Institute of Radiology Center for High Performance Computing and Washington University Research Infrastructure Services for use of their cluster computing platforms. The Artyomov lab was supported by grant from Aging Biology Foundation.

## Author Contributions

A.L.K., G.B., and A.B. generated single cell sequencing libraries. A.L.K., I.S., P.S.A, K.Z., L.L., and J.A. performed the analysis of single cell and single nucleus sequencing data. A.L.K., L.L., and G.S. performed the RNA in situ hybridization experiments. C.J., E.T., and S.L.R contributed to human cardiac tissue acquisition and processing. K.L. and M.A. are responsible for all aspects of this manuscript including experimental design, data analysis, and manuscript production.

## Statement on Human Specimens

This study complies with all relevant ethical regulations and was approved by the Washington University Institutional Review Board (Study #: 201104172). All samples were procured, and informed consent obtained by Washington University School of Medicine. BRISQ data can be found in Table S4.

## Methods

### Sample Preparation for Single Cell RNA Sequencing

Fresh cardiac tissues from LVAD cores or identical regions from the apex of explanted donors were minced with a razor blade and transferred into a 15ml conical tube containing DMEM with Collagenase I (450U/ml), DNAse I (60U/ml), and Hyaluronidase (60U/ml) and incubated at 37°C for 1 hour with agitation. Digestion was then stopped by addition of HBB buffer (2% FBS, 0.2% BSA in HBSS) and filtered through a 40um filter into 50 mL conical, transferred to a clean 15ml conical and centrifuged at 350 × G for 5 minutes at 4°C. Supernatant was then removed and pellet resuspended in 1mL ACK lysing buffer (Gibco A10492) and incubated at room temperature for 5 minutes followed by the addition of 9mL DMEM. Suspension was then centrifuged at above conditions, followed by removal of supernatant, and resuspension in 5mL FACS buffer (2% FBS, 2mM EDTA in calcium/magnesium free PBS). Centrifugation was repeated at above conditions, supernatant removed, and pellet resuspended in 300uL cell resuspension buffer (0.04% BSA in 1X PBS) and 1uL each of DRAQ5 (Thermo Scientific 62251) and DAPI (BD Biosciences 564907) and allowed to incubate 5 minutes before sorting. DRAQ5+/DAPI-cells were collected in cell resuspension buffer. Collected cells were then recentrifuged according to above parameters and resuspended in cell resuspension buffer to a target concentration of 1000 cells/μL. Nuclei were counted on a hemocytometer and concentration adjusted as necessary.

### Sample Preparation for Single Nuclei RNA Sequencing

Frozen cardiac tissues from LVAD cores or identical region from the apex of explanted donors were minced with a razor blade and transferred into a small (5 mL) Dounce homogenizer containing 1-2mL of chilled lysis buffer (10mM Tris-HCl pH 7.4, 10mM NaCl, 3mM MgCl_2_, 0.1% NP-40 in nuclease free water). Homogenized gently using 5 passes without rotation, then incubated on ice for 15 minutes. Lysate was then gently filtered through a 40um filter into 50mL conical, followed by rinsing the filter once with 1 mL lysis buffer, and transfer of lysate to a new 15ml conical tube. Nuclei were then centrifuged at 500 × g for 5 min at 4°C Followed by resuspension in 1mL Nuclei Wash Buffer (2% BSA, 0.2U/μL RNAse inhibitor in 1x PBS) and filtered through a 20 um pluristrainer into a fresh 15mL conical. Centrifugation was repeated according to above parameters, Supernatant was then removed and nuclei resuspended in 300uL Nuclei Wash Buffer and transferred to 5 mL tube for flow sorting. 1uL DRAQ5 (5 mM solution, Thermo Cat #62251) was added, mixed gently, and allowed to incubate 5 minutes before sorting. DRAQ5+ nuclei were sorted into Nuclei Wash Buffer on BD FACS Melody (BD Biosciences, San Jose, CA) using a 100 μM nozzle. Recovered nuclei were centrifuged again at the above parameters and were gently resuspended in Nuclei Wash Buffer to a target concentration of 1000 nuclei/uL. Nuclei were counted on a hemocytometer and concentration adjusted as necessary.

### Single Cell/Nuclei RNA Sequencing Analysis

Cells and Nuclei were processed using the Chromium Single Cell 5’ Reagent V1.1 Kit from 10X Genomics. 10,000 cells or nuclei per sample were loaded into a Chip G for GEM generation. Reverse transcription, barcoding, cDNA amplification, and purification for library preparation were performed according to the Chromium 5’ V1.1 protocol. Sequencing was performed on a NovaSeq 6000 platform (Illumina) targeting 100,000 reads/cell or nucleus. Cells were aligned to the human GRCh38 transcriptome while nuclei were aligned to the whole genome pre-MRNA reference generated from the GRCh38 transcriptome using the CellRanger V3 software (10X Genomics) according to the 10X Genomics instructions. Filtering, unsupervised clustering, differential expression, and additional analysis were completed using R and Python, including Seurat V3 and V4 and clusterProfiler packages for R and Palantir Python package.^11,39–41^

### QC, Filtering, and Clustering

For independent cell and nuclei analyses, individual sample matrices were imported into the Seurat v3.2.3 R package and combined into a Seurat object. Cells were filtered for mitochondrial reads <10%, and 2000<nCount_RNA<10000. Nuclei were filtered for mitochondrial reads <5%, and 1000<nCount_RNA<10000. No filtering was applied based on nFeature_RNA. The objects were then saved for easy import after manual doublet removal. For each object, transformation and normalization was performed using SCTransform to fit a negative binomial distribution and regress out mitochondrial read percentage. Principle components were then calculated (60 PCs for cells and 80 PCs for nuclei) and an elbow plot generated to select the cutoff for significant PCs to use for downstream analysis. UMAP dimensional reduction was then computed using the selected significant PCs (40 for cells and 80 for nuclei). Unsupervised clustering was then performed using the FindNeighbors and FindClusters function, again using the selected significant PC level as above, calculating clustering at a range of resolutions between 0.01-1. Differential gene expression was performed using the FindAllMarkers command using default parameters at high clustering resolution to aid in manual doublet discovery.

Doublets were visually identified on UMAP feature plots for nCount_RNA as well as feature plots for genes and gene z-scores enriched in each cluster. Clusters were considered to be doublets when nCount_RNA values were extraordinarily higher than other clusters and overlap of genes and z-scores for multiple independent populations. Small clusters that appeared between larger groups of clusters were especially scrutinized as these are more likely to exhibit doublets of cells from the clusters they exist between. After identification, these cells were removed, and the list of remaining cells saved. Raw objects from above were then loaded, subset to include cells that remained after doublet removal, and clustering repeated starting with transformation and normalization. This doublet removal process was repeated twice for the cell object and three times for the nuclei object until no doublet clusters were apparent. Final resolutions used for analysis were selected following detection of differentially expressed genes at multiple resolutions and identifying the highest resolution at which significantly enriched genes were still present in each cluster (final resolution used was 0.6 for cell object and 0.5 for nuclei object). Metadata for condition, age, sex, and cell type name were also added to the final objects.

### Integration of Single Cell and Single Nuclei Datasets

Integration of single cell and single nuclei datasets was performed using reference mapping in Seurat v4.0.1. A filtered and SCTransformed object from the single cell dataset but without calculation of PCs or clustering was loaded along with the final clustered single nuclei dataset object. Reference anchors between the reference nuclei and query cell datasets were identified using the SCT normalization and PCA reduction with 80 PCs from the nuclei object using the FindTransferAnchors command. The single cell object was then mapped to the single nuclei object using the MapQuery command using PCA and UMAP reference reduction. Mapping was performed at two levels (individual Seurat cluster and cell type) in order to visualize predictions scores at high and low granularity. The single cells and nuclei objects were then merged and a new UMAP reduction computed. Reclustering was then performed by utilizing the FindNeighbors and FindClusters command at multiple resolutions between 0.1-1. No doublet exclusion or filtering was necessary as mapping and integration was performed on already filtered objects. The final resolution was selected to be 0.6 as this resolution captured the distinct clusters found in both single cell and single nuclei analysis. Metadata for condition, age, sex, and cell type name were also added to the final object.

### Detection of Differentially Expressed Genes (DEGs)

Detection of differentially expressed genes between clusters was performed using the FindAllMarkers command, specifying return of only upregulated genes with a Log_2_FC cutoff of 0.1. For downstream analysis, DEGs were further filtered by Log_2_FC and p-value as described for that analysis. For individual cell types, differential expression comparing only 2 groups by condition, sex, or age was performed using the FindMarkers function specifying no minimum percentage of cells expressing an individual gene, return of both positive and negatively changed genes, and no cutoffs for Log_2_FC or p-value in order to obtain even non-significant changes in expression for every gene present in the analysis. Filtering of this DEG table was performed by Log_2_FC and p-value for further analysis as described in the manuscript. For all DEG calculations the default ‘SCT’ assay and ‘data’ slot were used and performed using the default Wilcoxan Rank Sum method

### Calculation of Population Z-Scores

Z-score values were calculated using R v3.6.2 and v4.0.1. For each population where z-scores were calculated, genesets used were selected based on high enrichment in a population based on DEG analysis described above. The expression matrix used to calculate Z-scores was extracted from a Seurat object using the GetAssayData function from the Seurat package from the default ‘SCT’ assay and ‘data’ slot. Z-scores were then calculated for each geneset for each individual cell or nuclei in the dataset by scaling gene expression within the matrix, setting NA values to 0 and using the following formula:

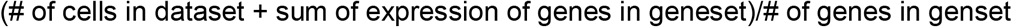

These calculated z-scores were appended to a table to be saved as well as each z-score added as metadata to the Seurat object for use in making feature plots.

### Pseudobulk RNA-seq

Pseduobulk RNA-seq analysis was performed using the DESeq2 package for R. A gene expression matrix was extracted from the Seurat object using the GetAssayData Seurat function specifying the ‘RNA’ assay and ‘counts’ slot to extract raw sequencing counts for each gene and cell. Counts in this matrix were then summed per gene for each sample into a new matrix. The resulting matrix was then normalized using DESeq2 by estimating size factors and performing normalization with the counts function resulting in a new matrix with normalized counts for each gene and sample similar to the output of a traditional bulk sequencing experiment. The DESeq function was then utilized to calculate differential gene expression based on negative binomial distribution. Pairwise comparisons were then completed by condition of interest (disease state, sex, age group) using the Wald test and an alpha value of 0.5 for independent filtering and adding Log_2_FC using the lfcShrink function with ‘ashr’ adaptive shrinking. We specified no cutoffs for Log_2_FC or p-value in order to obtain even non-significant changes in expression. Filtering of this DEG table was performed by LogFC and p-value for further analysis as described in the manuscript.

### Pathway Analysis

Pathway analysis was completed using the ClusterProfiler R package. A list of genes present in both the Seurat and Pseudobulk differential expression analyses by disease state with Log_2_FC>0.1 and adjusted p-value<0.05 was utilized in the pathway analysis. Genes with negative and positive Log_2_FC values were separated in order to identify enrichment in either the non-diseased or diseased condition, respectively. The enrichWP function was used to return a table with pathway enrichments from the WikiPathways database.

For comparison of enriched pathways between multiple populations/states, the compareCluster function was utilized on a matrix from the output Seurat differential expression analysis filtered for Log_2_FC>0.1 and adjusted p-value<0.05 that contained the column specifying in which population/state the gene was upregulated. This analysis utilized the enrichPathway database from ClusterProfiler to return a table of enriched pathways in each population/state.

### Transcription Factor Analysis

Transcription factor analysis was performed using the Enrichr web utility (https://maayanlab.cloud/Enrichr/enrich). Genes upregulated in a population/state based on Seurat differential expression analysis filtered for Log_2_FC>0.1 and adjusted p-value<0.05 were entered into the Enrichr and results from enrichment in the ChEA 2016 ChIP-seq database were downloaded and loaded as a matrix in R v4.0.1 for generation of dot plots.

### Trajectory Analysis

Trajectory analysis was performed using the Palantir package for Python. Using the normalized and scaled gene counts for the 3,000 highly variable genes a matrix was exported as the input. Using the matrix, principal components were calculated and then diffusion maps were calculated as an estimate of the low dimensional phenotypic manifold of the data. Then, the actual Palantir was run by specifying a start cell state (the progenitor cell type from the dataset). Palantir then returned the terminal cell states, entropy values, pseudotime values, and the probability of ending up in each of the terminal states for all cells.

### RNAScope in-situ hybridization

RNA was visualized using RNAScope Multiplex Fluorescent Reagent Kit v2 Assay, RNAScope 2.5 HD Detection Reagent – RED, and RNAScope 2.5 HD Duplex Assay kits (Advanced Cell Diagnostics) using probes designed by Advanced Cell Diagnostics for ANKRD1, MYH6, NPPA, NPPB, CD163, DCN, POSTN, PLA2G2A, CCL2, PCOLCE2, ELN, and RGS5.^42^ Samples were fixed for 24 hours at 4°C in 10% neutral buffered formalin. Samples were then washed in 1X PBS, equilibrated in 30% sucrose and embedded in OCT medium (Sakura Finetek) and stored at −80°C (fluorescence) or washed in 1X PBS, dehydrated in ethanol and embedded in paraffin (red and duplex). OCT embedded sections were cut at 12μm and paraffin embedded sections cut at 8μm. Fluorescent images were collected using a Zeiss LSM 700 laser scanning confocal microscope. Chromogenic/brightfield images were acquired using a Zeiss Axioscan Z1 automated slide scanner. Image processing was performed using Zen Blue and Zen Black (Zeiss), FIJI/ImageJ^43,44^, and Photoshop (Adobe).

## Data and Code Availability

The processed single cell objects that support the findings of this study are available on the Gene Expression Omnibus (GEO, ACCESION NUMBER TO BE ADDED PRIOR TO PUBLICATION). Scripts and methods used in processing can be found at https://github.com/alkoenig/Atlas_of_Human_Heart_Failure_Lavine. Sequences and raw expression matrices available from the authors upon request.

