## Supplementary Material for "Single Cell Transcriptomics Reveals Cell Type Specific Diversification in Human Heart Failure"

| | HEALTHY (N=26) | DCM (N=12) |
|---|---|---|
| **Demographics** | | |
| Mean age (years) | 49.4 ± 19.0 | 45.1 ± 19.8 |
| Median age (years) | 50.5 | 53.5 |
| Age range (years) | 11 – 75 | 8 – 64 |
| Sex – n. (%) | | |
| Male | 12 (46.2) | 7 (58.3) |
| Female | 14 (53.8) | 5 (41.7) |
| Race – n. (%) | | |
| White | 19 (73.1) | 9 (75.0) |
| Black or African American | 7 (27.9) | 3 (25.0) |
| Ethnicity – n. (%) | | |
| Non-Hispanic | 26 (100) | 11 (91.7) |
| Hispanic | 0 (0) | 1 (8.3) |
| **Baseline Clinical Medical History** | | |
| BMI (kg/m$^2$) – mean ± SD | 28.8 ± 7.9 | 26.7 ± 6.2 |
| Hypertension – n. (%) | 14 (53.9) | 4 (33.3) |
| Diabetes – no. (%) | 9 (34.6) | 4 (33.3) |
| Chronic Kidney Disease – n. (%) | 2 (7.7) | 4 (33.3) |
| Smoking – n. (%) | 15 (57.7) | 5 (41.7) |
| **Cardiac Clinical Data** | | |
| Cardiac Ejection Fraction – mean ± SD | 60.7 ± 7.8 | 19.8 ± 6.9 |
| Cardiac Output (L/min) – mean ± SD | 7.9 ± 2.5 | 2.7 ± 0.4 |
| Arrhythmias – n. (%) | 4 (15.4) | 9 (75.0) |
| Valve Disease (≥mild) – n. (%) | 8 (30.8) | 9 (75.0) |
| Mitral valve | 6 (23.1) | 9 (75.0) |
| Aortic valve | 3 (11.4) | 1 (8.3) |
| Pulmonary valve | 1 (3.9) | 1 (8.3) |
| Tricuspid valve | 5 (19.2) | 5 (41.7) |

**Table S1. Demographic and Baseline Clinical Data Table for Single-Nuclei RNA Sequencing Analysis**

| | HEALTHY (N=2) | DCM (N=6) |
|---|---|---|
| **Demographics** | | |
| Mean age (years) | 63 ± 0 | 45.7 ± 20.7 |
| Median age (years) | 63 | 46.5 |
| Age range (years) | 63 – 63 | 23 – 74 |
| Sex – n. (%) | | |
| Male | 1 (50.0) | 5 (83.3) |
| Female | 1 (50.0) | 1 (16.7) |
| Race – n. (%) | | |
| White | 1 (50.0) | 3 (50.0) |
| Black or African American | 1 (50.0) | 3 (50.0) |
| Ethnicity – n. (%) | | |
| Non-Hispanic | 2 (100) | 6 (100) |
| Hispanic | 0 (0) | 0 (0) |
| **Baseline Clinical Medical History** | | |
| BMI ($kg/m^2$) – mean ± SD | 27.8 ± 0.5 | 26. 8 ± 9.7 |
| Hypertension – n. (%) | 2 (100) | 4 (66.7) |
| Diabetes – no. (%) | 0 (0) | 2 (33.3) |
| Chronic Kidney Disease – n. (%) | 0 (0) | 3 (50.0) |
| Smoking – n. (%) | 2 (100) | 5 (83.3) |
| **Cardiac Clinical Data** | | |
| Cardiac Ejection Fraction – mean ± SD | 56.5 ± 9.2 | 20.7 ± 5.0 |
| Cardiac Output (L/min) – mean ± SD | 6.5 ± 0.1 | 4.1 ± 1.9 |
| Arrhythmias – n. (%) | 0 (0) | 4 (66.7) |
| Valve Disease (≥mild) – n. (%) | 0 (0) | 5 (83.3) |
| Mitral valve | 0 (0) | 4 (66.7) |
| Aortic valve | 0 (0) | 1 (16.7) |
| Pulmonary valve | 0 (0) | 2 (33.3) |
| Tricuspid valve | 0 (0) | 3 (50.0) |

**Table S2. Demographic and Baseline Clinical Data Table for Single-Cell RNA Sequencing Analysis**

| | HEALTHY (N=28) | DCM (N=18) |
|---|---|---|
| **Demographics** | | |
| Mean age (years) | 50.2 ± 18.2 | 45.3 ± 19.5 |
| Median age (years) | 53 | |
| Age range (years) | 11 – 75 | 8 – 74 |
| Sex – n. (%) | | |
| Male | 13 (46.4) | 12 (66.7) |
| Female | 15 (53.6) | 6 (21.4) |
| Race – n. (%) | | |
| White | 20 (71.4) | 12 (66.7) |
| Black or African American | 8 (28.6) | 6 (21.4) |
| Ethnicity – n. (%) | | |
| Non-Hispanic | 28 (100) | 17 (94.4) |
| Hispanic | 0 (0) | 1 (5.6) |
| **Baseline Clinical Medical History** | | |
| BMI (kg/m$^2$) – mean ± SD | 28.8 ± 7.4 | 26.8 ± 7.2 |
| Hypertension – n. (%) | 16 (57.1) | 8 (44.4) |
| Diabetes – no. (%) | 9 (32.1) | 6 (33.3) |
| Chronic Kidney Disease – n. (%) | 2 (7.1) | 7 (38.9) |
| Smoking – n. (%) | 17 (60.7) | 9 (50.0) |
| **Cardiac Clinical Data** | | |
| Cardiac Ejection Fraction – mean ± SD | 58.7 ± 11.1 | 19.8 ± 6.9 |
| Cardiac Output (L/min) – mean ± SD | 7.7 ± 2.4 | 3.2 ± 1.3 |
| Arrhythmias – n. (%) | 4 (14.3) | 13 (72.2) |
| Valve Disease (≥mild) – n. (%) | 8 (28.6) | 14 (77.8) |
| Mitral valve | 6 (21.4) | 13 (72.2) |
| Aortic valve | 3 (10.7) | 2 (11.1) |
| Pulmonary valve | 1 (3.6) | 3 (16.7) |
| Tricuspid valve | 5 (17.9) | 8 (44.4) |

**Table S3. Demographic and Baseline Clinical Data Table for Combined Single-Nuclei and Single-Cell RNA Sequencing Analysis**

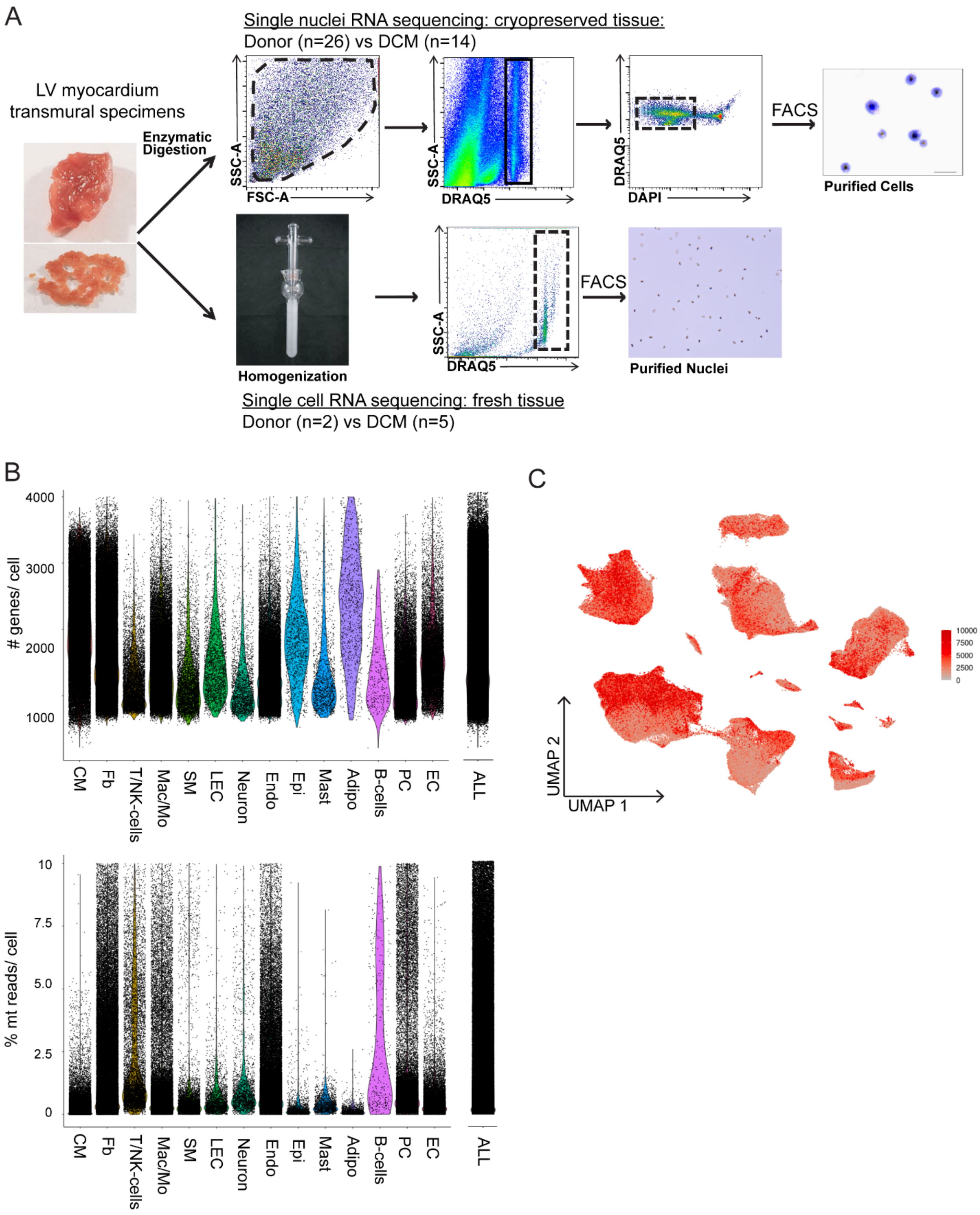
A
Single nuclei RNA sequencing: cryopreserved tissue:
Donor (n=26) vs DCM (n=14)
LV myocardium
transmural specimens
Enzymatic
Digestion
SSC-A
FSC-A
DRAQ5
DAPI
FACS
Purified Cells
Homogenization
Purified Nuclei
Single cell RNA sequencing: fresh tissue
Donor (n=2) vs DCM (n=5)
B
### genes/ cell
4000
3000
2000
1000
CM
Fb
T/NK-cells
Mac/Mo
SM
LEC
Neuron
Endo
Epi
Mast
Adipo
B-cells
PC
EC
ALL
% mt reads/ cell
10
7.5
5.0
2.5
0
C
UMAP 2
UMAP 1
10000
7500
5000
2500
0

**Figure S1. Sample processing and QC Plots**

**A.** Diagram of tissue processing and flow cytometry cell sorting strategies for single nucleus RNA sequencing (top) and single cell RNA sequencing (bottom). **B.** Violin plots of the number of genes (top) and percent mitochondrial reads (bottom) per cell/nuclei split by cell type for the integrated Seurat object after QC filtering. **C.** UMAP feature plot of number of reads (UMI) per cell/nuclei from integrated Seurat object after QC filtering.

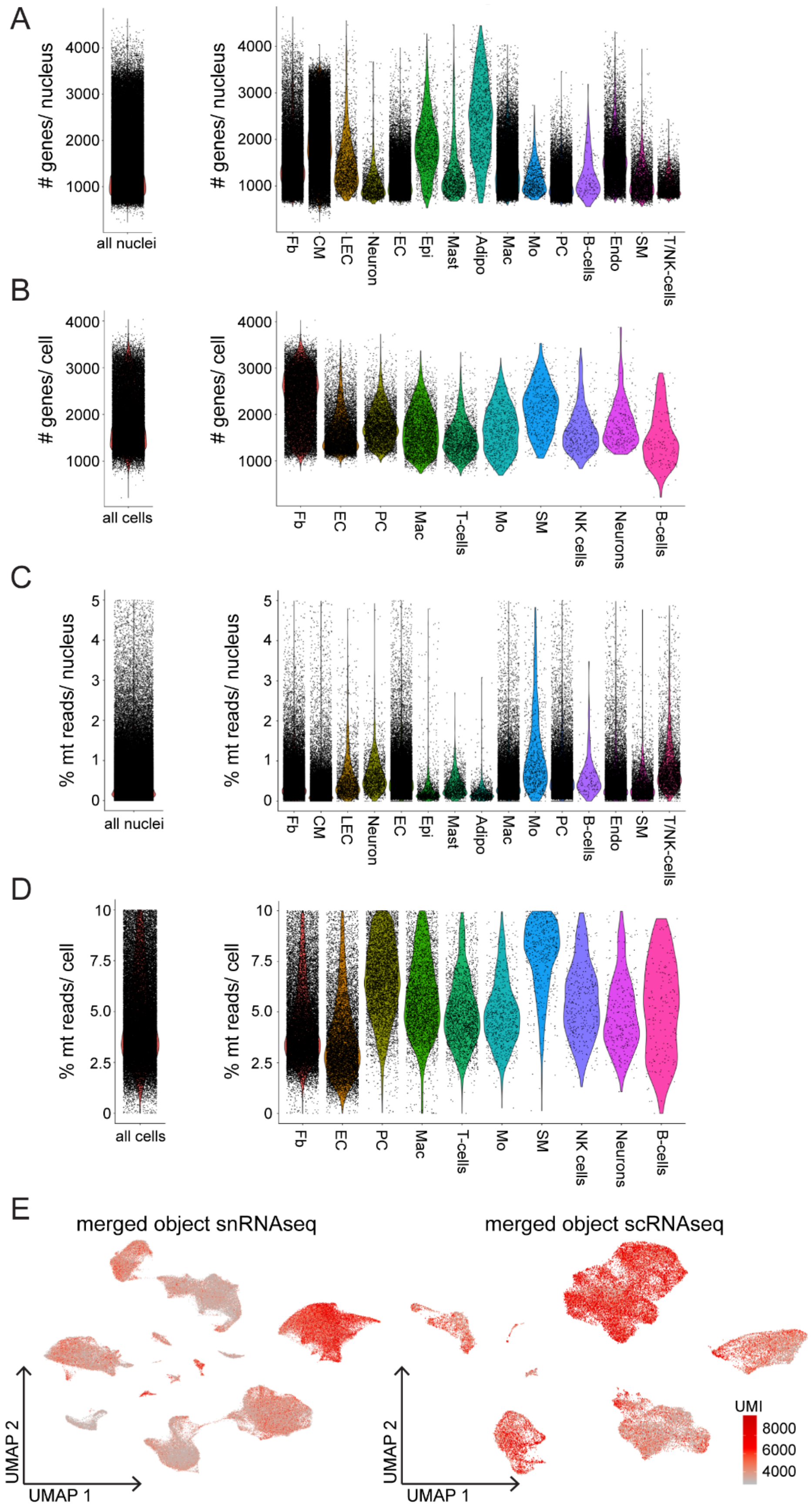

A
### genes/ nucleus
4000
3000
2000
1000
all nuclei
Fb
CM
LEC
Neuron
EC
Epi
Mast
Adipo
Mac
Mo
PC
B-cells
Endo
SM
T/NK-cells
B
### genes/ cell
all cells
Fb
EC
PC
Mac
T-cells
Mo
SM
NK cells
Neurons
B-cells
C
% mt reads/ nucleus
5
4
3
2
1
0
D
% mt reads/ cell
10
7.5
5.0
2.5
0
E
merged object snRNAseq
merged object scRNAseq
UMAP 2
UMAP 1
UMI
8000
6000
4000

**Figure S2. QC Plots for individual snRNA seq and scRNA seq objects**

**A.** Violin plot of the number of genes per nucleus from the single nucleus RNA sequencing dataset after QC filtering split by cell type. **B.** Violin plot of number of genes per cell from the single cell RNA sequencing dataset after QC filtering split by cell type. **C.** Violin plot of percent mitochondrial reads per nucleus from the single nucleus RNA sequencing dataset after QC filtering split by cell type. **D.** Violin plot of percent mitochondrial reads per cell from the single cell RNA sequencing dataset after QC filtering split by cell type. **E.** UMAP feature plots of number of reads (UMI) per cell/nuclei from single cell RNA sequencing (left) and single nucleus RNA sequencing (right) datasets after QC filtering.

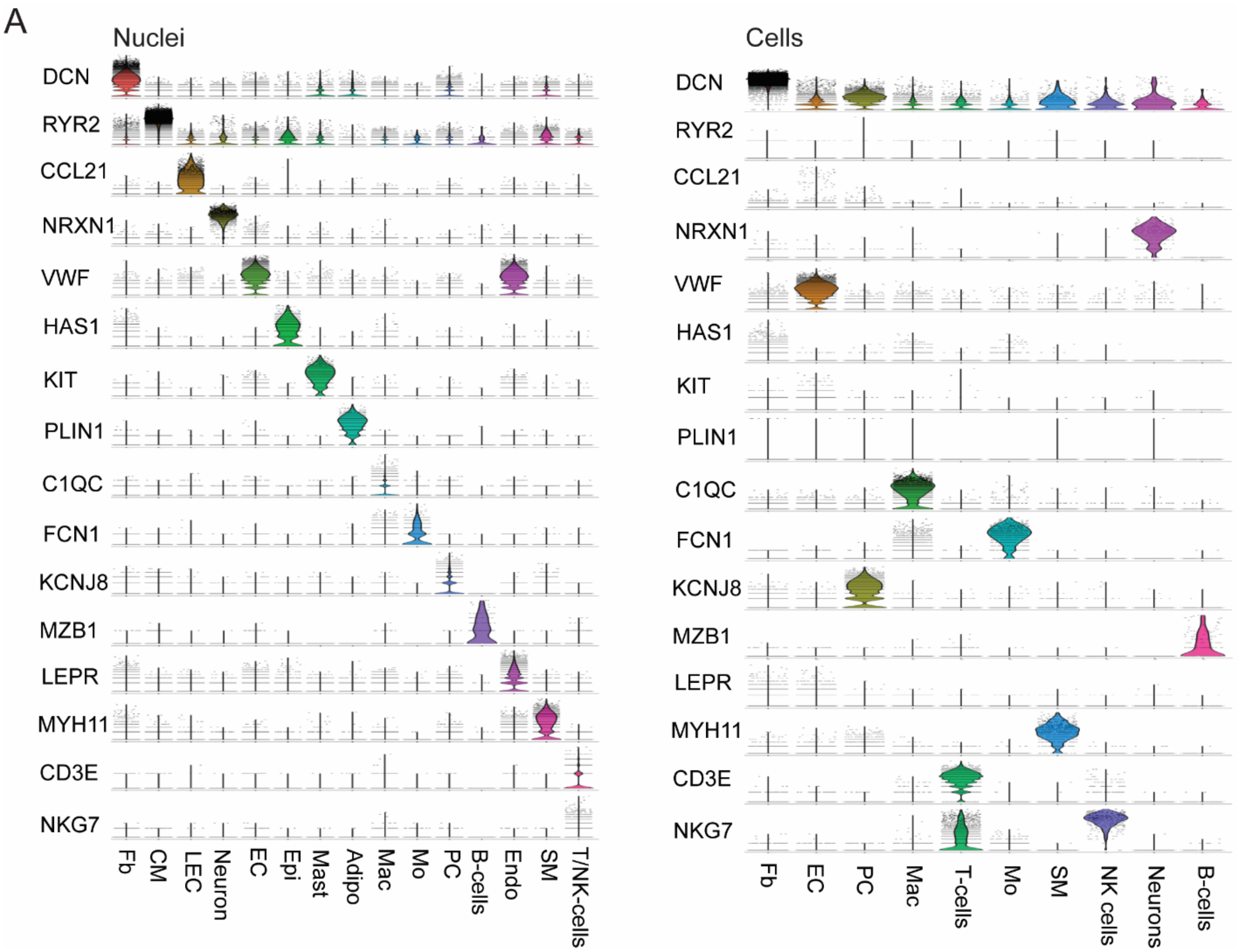
A
Nuclei
DCN
RYR2
CCL21
NRXN1
VWF
HAS1
KIT
PLIN1
C1QC
FCN1
KCNJ8
MZB1
LEPR
MYH11
CD3E
NKG7
Fb
CM
LEC
Neuron
EC
Epi
Mast
Adipo
Mac
Mo
PC
B-cells
Endo
SM
T/NK-cells
Cells
Fb
EC
PC
Mac
T-cells
Mo
SM
NK cells
Neurons
B-cells

**Figure S3.**

Violin plots split by cluster displaying the expression of characteristic cell marker genes in the single nucleus RNA sequencing (left) and single cell RNA sequencing (right) datasets.

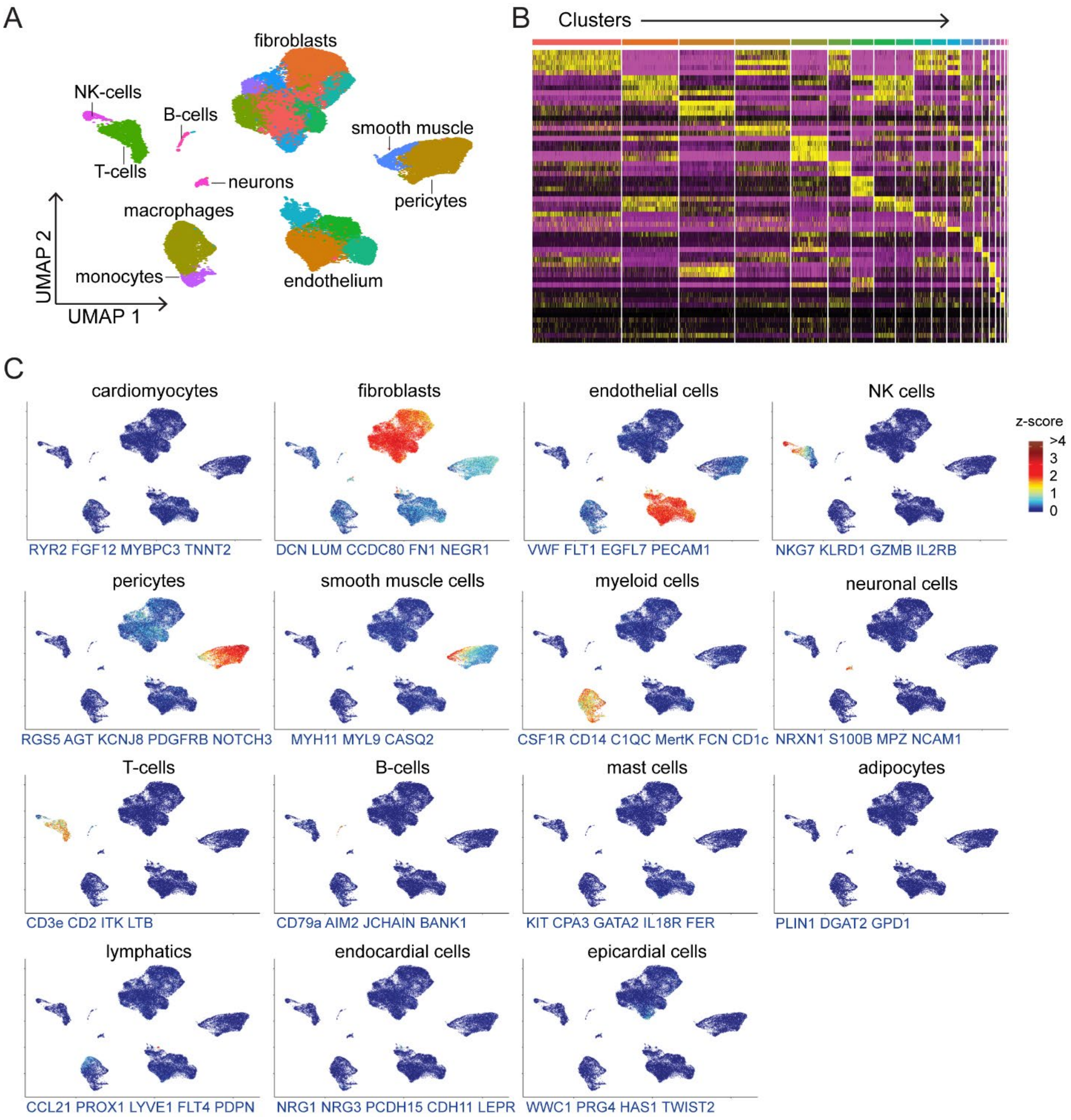
A
fibroblasts
NK-cells
B-cells
smooth muscle
T-cells
neurons
pericytes
macrophages
UMAP 2
monocytes
endothelium
UMAP 1
B
Clusters
C
cardiomyocytes
RYR2 FGF12 MYBPC3 TNNT2
fibroblasts
DCN LUM CCDC80 FN1 NEGR1
endothelial cells
VWF FLT1 EGFL7 PECAM1
NK cells
NKG7 KLRD1 GZMB IL2RB
z-score
>4
3
2
1
0
pericytes
RGS5 AGT KCNJ8 PDGFRB NOTCH3
smooth muscle cells
MYH11 MYL9 CASQ2
myeloid cells
CSF1R CD14 C1QC MertK FCN CD1c
neuronal cells
NRXN1 S100B MPZ NCAM1
T-cells
CD3e CD2 ITK LTB
B-cells
CD79a AIM2 JCHAIN BANK1
mast cells
KIT CPA3 GATA2 IL18R FER
adipocytes
PLIN1 DGAT2 GPD1
lymphatics
CCL21 PROX1 LYVE1 FLT4 PDPN
endocardial cells
NRG1 NRG3 PCDH15 CDH11 LEPR
epicardial cells
WWC1 PRG4 HAS1 TWIST2

**Figure S4. Single cell RNA sequencing identifies 10 cell types within the LV myocardium.**

**A.** UMAP projection showing unsupervised clustering of single cell RNA sequencing data. **B.** Heatmap of the top 10 genes by log2FC enriched in each cluster. **C.** Z-score feature plots for transcriptional signatures enriched in each cell type. Genes used for cell type identification (blue) were selected based on enrichment from Seurat differential expression analysis.

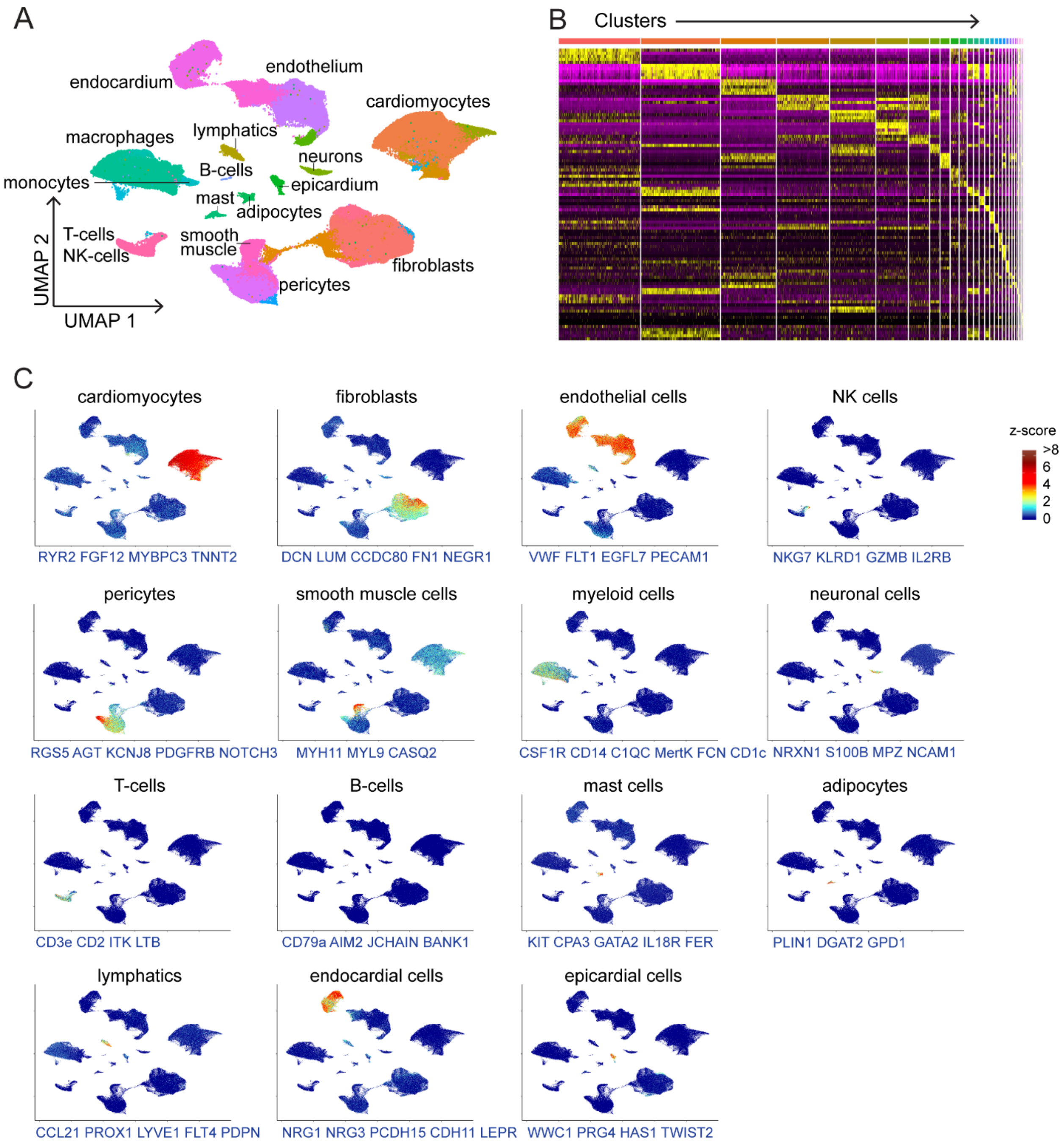

A
endocardium
endothelium
cardiomyocytes
macrophages
lymphatics
neurons
monocytes
B-cells
epicardium
mast
adipocytes
T-cells
NK-cells
smooth
muscle
fibroblasts
pericytes
UMAP 2
UMAP 1
B
Clusters
C
cardiomyocytes
RYR2 FGF12 MYBPC3 TNNT2
fibroblasts
DCN LUM CCDC80 FN1 NEGR1
endothelial cells
VWF FLT1 EGFL7 PECAM1
NK cells
NKG7 KLRD1 GZMB IL2RB
z-score
>8
6
4
2
0
pericytes
RGS5 AGT KCNJ8 PDGFRB NOTCH3
smooth muscle cells
MYH11 MYL9 CASQ2
myeloid cells
CSF1R CD14 C1QC MertK FCN CD1c
neuronal cells
NRXN1 S100B MPZ NCAM1
T-cells
CD3e CD2 ITK LTB
B-cells
CD79a AIM2 JCHAIN BANK1
mast cells
KIT CPA3 GATA2 IL18R FER
adipocytes
PLIN1 DGAT2 GPD1
lymphatics
CCL21 PROX1 LYVE1 FLT4 PDPN
endocardial cells
NRG1 NRG3 PCDH15 CDH11 LEPR
epicardial cells
WWC1 PRG4 HAS1 TWIST2

**Figure S5. Single nucleus RNA sequencing identifies additional cell populations within the LV myocardium not recovered by single cell RNA sequencing.**

**A.** UMAP projection showing unsupervised clustering of single nucleus RNA sequencing data. **B.** Heatmap of the top 10 genes by log2FC enriched in each cluster. **C.** Z-score feature plots for transcriptional signatures enriched in each cell type. Genes used for cell type identification (blue) were selected based on enrichment from Seurat differential expression analysis.

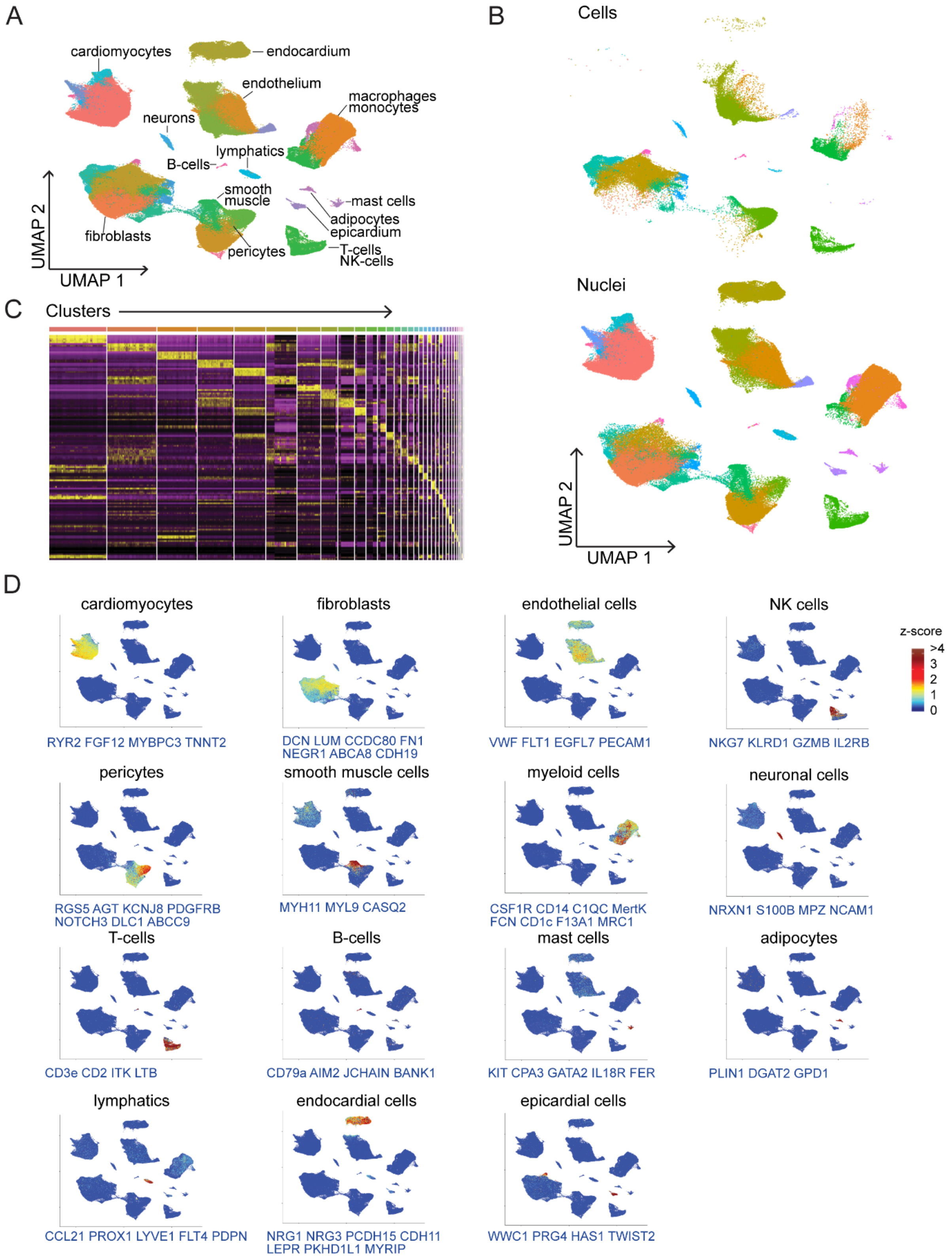

A
cardiomyocytes
endocardium
endothelium
macrophages
monocytes
neurons
lymphatics
B-cells
smooth muscle
mast cells
adipocytes
epicardium
fibroblasts
pericytes
T-cells
NK-cells
UMAP 2
UMAP 1
B
Cells
Nuclei
UMAP 2
UMAP 1
C
Clusters
D
cardiomyocytes
RYR2 FGF12 MYBPC3 TNNT2
fibroblasts
DCN LUM CCDC80 FN1 NEGR1 ABCA8 CDH19
endothelial cells
VWF FLT1 EGFL7 PECAM1
NK cells
NKG7 KLRD1 GZMB IL2RB
z-score
>4
3
2
1
0
pericytes
RGS5 AGT KCNJ8 PDGFRB NOTCH3 DLC1 ABCC9
smooth muscle cells
MYH11 MYL9 CASQ2
myeloid cells
CSF1R CD14 C1QC MertK FCN CD1c F13A1 MRC1
neuronal cells
NRXN1 S100B MPZ NCAM1
T-cells
CD3e CD2 ITK LTB
B-cells
CD79a AIM2 JCHAIN BANK1
mast cells
KIT CPA3 GATA2 IL18R FER
adipocytes
PLIN1 DGAT2 GPD1
lymphatics
CCL21 PROX1 LYVE1 FLT4 PDPN
endocardial cells
NRG1 NRG3 PCDH15 CDH11 LEPR PKHD1L1 MYRIP
epicardial cells
WWC1 PRG4 HAS1 TWIST2

**Figure S6. Integration of single cell RNA sequencing and single nucleus RNA sequencing data allows for combined analysis of samples from different technologies.**

**A.** UMAP projection showing unsupervised clustering of the integrated dataset. **B.** UMAP projection split by technology. **C.** Heatmap of the top 10 genes by log2FC enriched in each cluster. **D.** Z-score feature plots for transcriptional signatures enriched in each cell type. Genes used for cell type identification (blue) were selected based on enrichment from Seurat differential expression analysis.

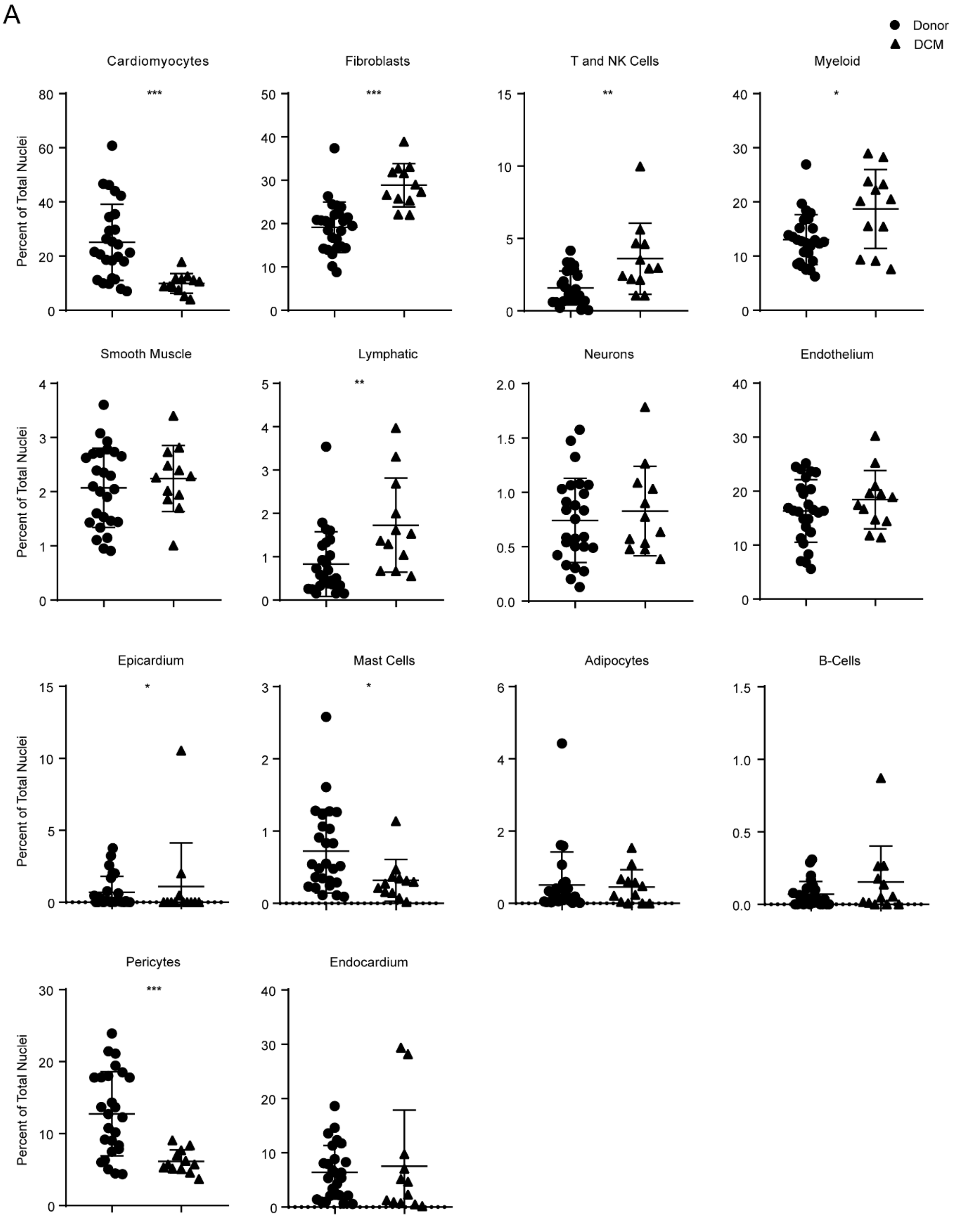
A
Donor
DCM
Cardiomyocytes
***
Fibroblasts
***
T and NK Cells
**
Myeloid
*
Smooth Muscle
Lymphatic
**
Neurons
Endothelium
Epicardium
*
Mast Cells
*
Adipocytes
B-Cells
Pericytes
***
Endocardium
Percent of Total Nuclei

**Figure S7. Distribution of nuclei by cluster identifies the relative distribution of myocardial cell types in donor controls and dilated cardiomyopathy.**

Dot plots displaying percent of total nuclei per sample assigned to each cell type based on unsupervised clustering in the integrated dataset. Each data point represents a subject. Lines and error bars are representative of mean and standard deviation, respectively. * $p<0.05$, ** $p<0.01$, *** $p<0.001$ using Mann-Whitney-U test, lines represent mean and standard deviation.

A

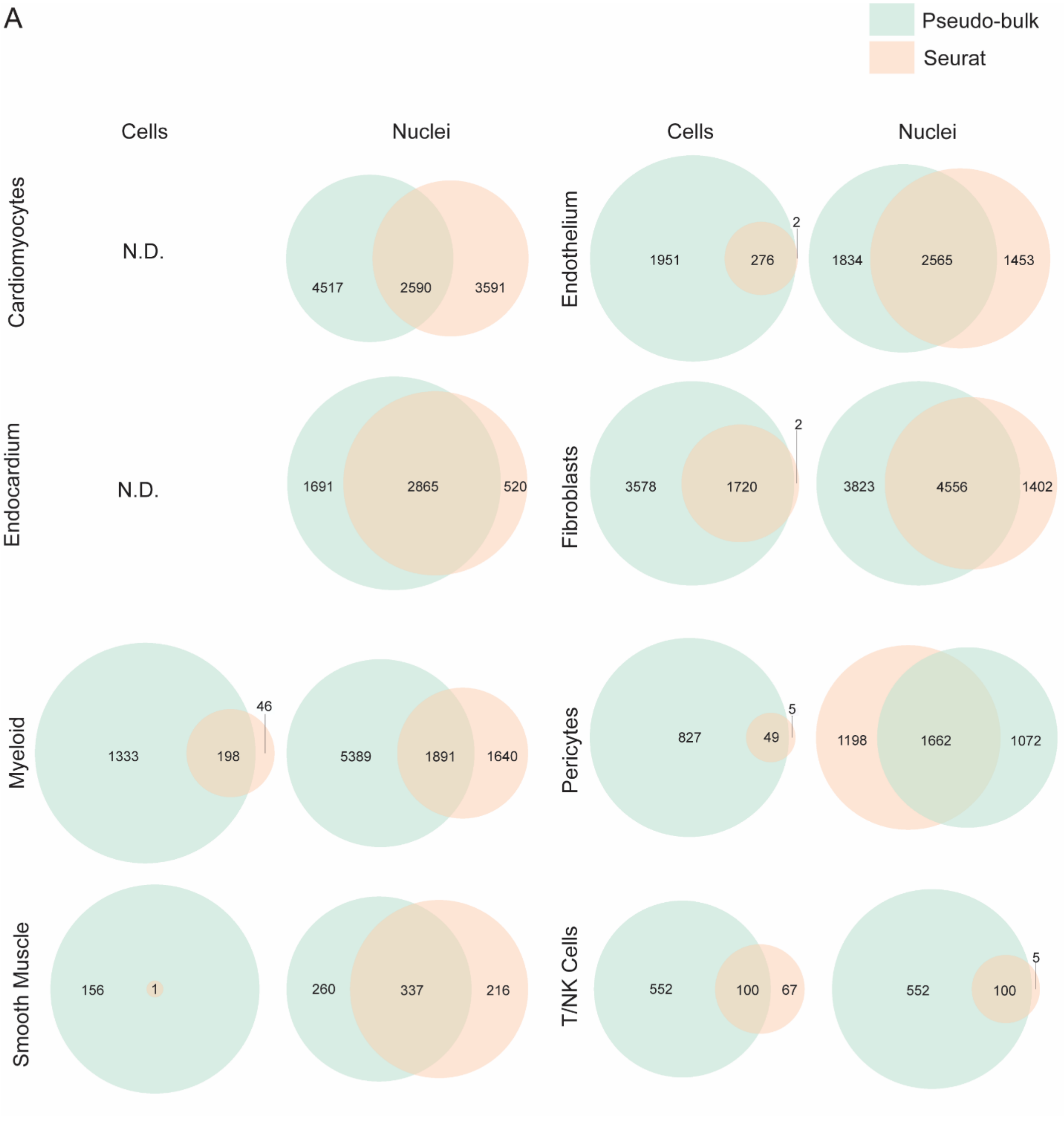
Pseudo-bulk
Seurat
Cells
Nuclei
Cells
Nuclei
Cardiomyocytes
N.D.
4517
2590
3591
Endothelium
1951
276
2
1834
2565
1453
Endocardium
N.D.
1691
2865
520
Fibroblasts
3578
1720
2
3823
4556
1402
Myeloid
1333
198
46
5389
1891
1640
Pericytes
827
49
5
1198
1662
1072
Smooth Muscle
156
1
260
337
216
T/NK Cells
552
100
67
552
100
5

**Figure S8. Pseudobulk and Seurat differential expression analyses identify overlapping sets of differentially regulated genes.**

Venn diagrams of differentially regulated genes (adjusted $p<0.05$) in each major cell type highlighting genes identified in the Seurat single cell differential expression analysis, DESeq2 pseudobulk differential expression analysis, and intersection of the Seurat and pseudobulk differential expression analyses (overlap). N.D. – Cell types not present in single cell RNA sequencing dataset.

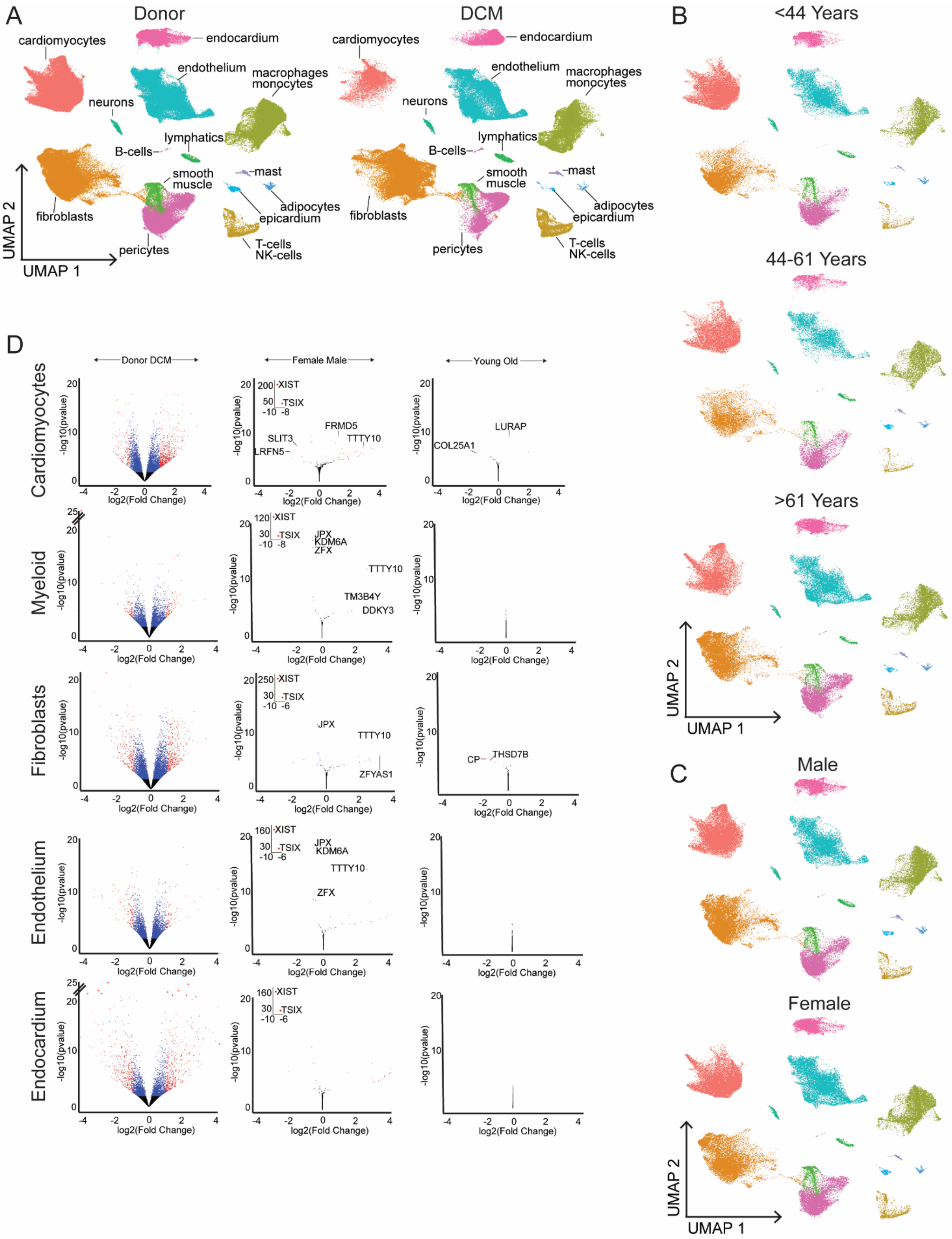
A
Donor
DCM
cardiomyocytes
endocardium
endothelium
macrophages
monocytes
neurons
lymphatics
B-cells
smooth
muscle
mast
adipocytes
epicardium
fibroblasts
pericytes
T-cells
NK-cells
UMAP 2
UMAP 1
B
<44 Years
44-61 Years
>61 Years
C
Male
Female
D
Donor DCM
Female Male
Young Old
Cardiomyocytes
Myeloid
Fibroblasts
Endothelium
Endocardium
-log10(pvalue)
log2(Fold Change)
XIST
TSIX
FRMD5
SLIT3
TTTY10
LRFN5
LURAP
COL25A1
JPX
KDM6A
ZFX
TM3B4Y
DDKY3
ZFYAS1
CP
THSD7B

**Figure S9. Pseudobulk differential expression reveals the contribution of disease state, sex, and age across major cell types.**

**A,** UMAP plot of complete integrated dataset split by condition and colored by cell type. **B.** UMAP plot of non-diseased donor samples from integrated dataset split by age tertile and colored by cell type. **C.** UMAP plot of non-diseased donor samples from integrated dataset split by sex and colored by cell type. **D.** Volcano plots of pseudobulk differential expression analysis of single nucleus RNA sequencing data performed on each cell type comparing donor control vs. dilated cardiomyopathy (DCM) (left), sex (center), and age (right). Younger (<52 years) and older groups (>52 years) were split based on the median age. Differential expression comparing sex and age group was performed on non-diseased donor samples.

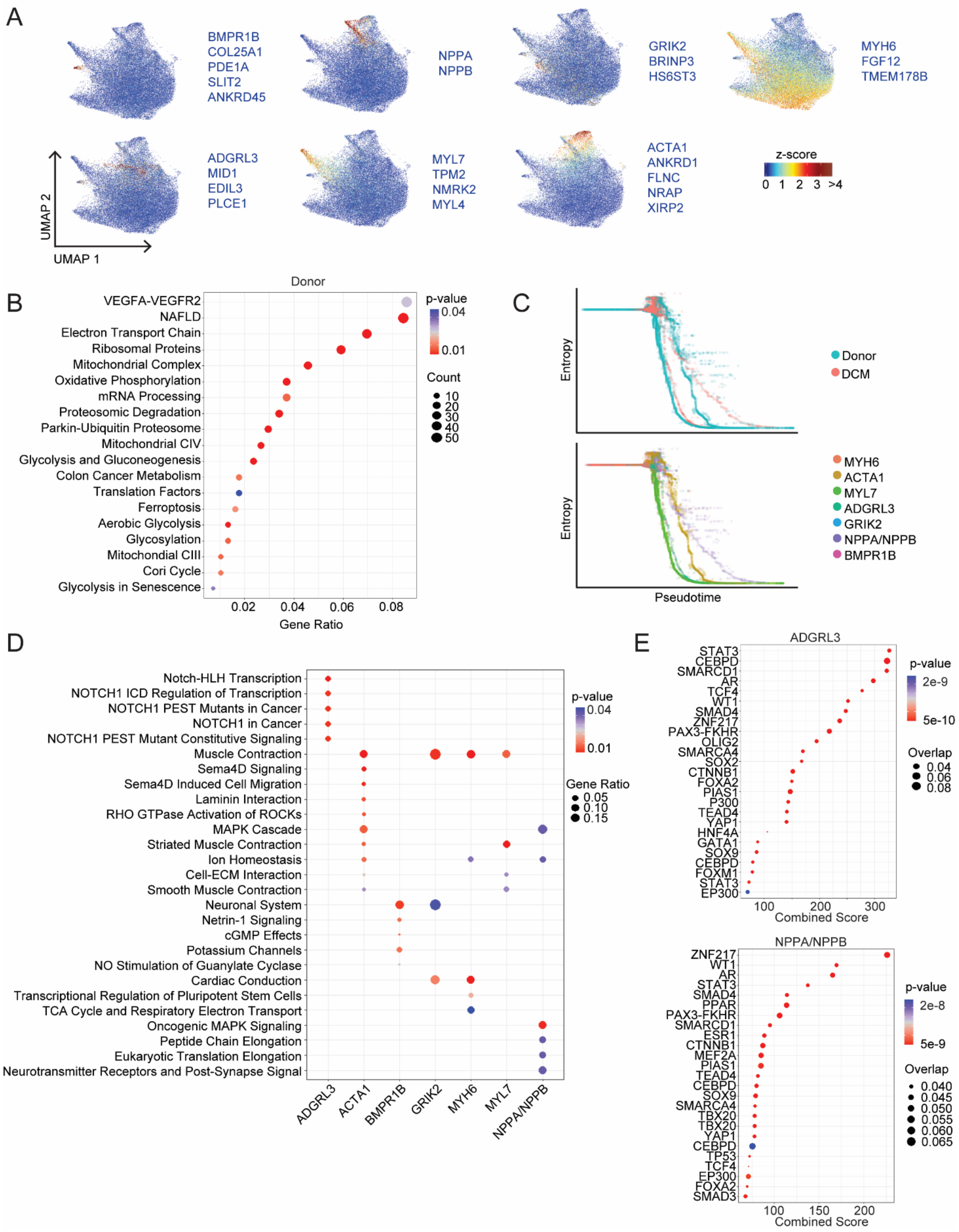
A
BMPR1B
COL25A1
PDE1A
SLIT2
ANKRD45
NPPA
NPPB
GRIK2
BRINP3
HS6ST3
MYH6
FGF12
TMEM178B
ADGRL3
MID1
EDIL3
PLCE1
MYL7
TPM2
NMRK2
MYL4
ACTA1
ANKRD1
FLNC
NRAP
XIRP2
z-score
0 1 2 3 >4
UMAP 2
UMAP 1
B
Donor
VEGFA-VEGFR2
NAFLD
Electron Transport Chain
Ribosomal Proteins
Mitochondrial Complex
Oxidative Phosphorylation
mRNA Processing
Proteosomic Degradation
Parkin-Ubiquitin Proteosome
Mitochondrial CIV
Glycolysis and Gluconeogenesis
Colon Cancer Metabolism
Translation Factors
Ferroptosis
Aerobic Glycolysis
Glycosylation
Mitochondrial CIII
Cori Cycle
Glycolysis in Senescence
p-value
0.04
0.01
Count
10
20
30
40
50
0.02 0.04 0.06 0.08
Gene Ratio
C
Entropy
Donor
DCM
MYH6
ACTA1
MYL7
ADGRL3
GRIK2
NPPA/NPPB
BMPR1B
Pseudotime
D
Notch-HLH Transcription
NOTCH1 ICD Regulation of Transcription
NOTCH1 PEST Mutants in Cancer
NOTCH1 in Cancer
NOTCH1 PEST Mutant Constitutive Signaling
Muscle Contraction
Sema4D Signaling
Sema4D Induced Cell Migration
Laminin Interaction
RHO GTPase Activation of ROCKs
MAPK Cascade
Striated Muscle Contraction
Ion Homeostasis
Cell-ECM Interaction
Smooth Muscle Contraction
Neuronal System
Netrin-1 Signaling
cGMP Effects
Potassium Channels
NO Stimulation of Guanylate Cyclase
Cardiac Conduction
Transcriptional Regulation of Pluripotent Stem Cells
TCA Cycle and Respiratory Electron Transport
Oncogenic MAPK Signaling
Peptide Chain Elongation
Eukaryotic Translation Elongation
Neurotransmitter Receptors and Post-Synapse Signal
ADGRL3
ACTA1
BMPR1B
GRIK2
MYH6
MYL7
NPPA/NPPB
p-value
0.04
0.01
Gene Ratio
0.05
0.10
0.15
E
ADGRL3
STAT3
CEBPD
SMARCD1
AR
TCF4
WT1
SMAD4
ZNF217
PAX3-FKHR
OLIG2
SMARCA4
SOX2
CTNNB1
FOXA2
PIAS1
P300
TEAD4
YAP1
HNF4A
GATA1
SOX9
CEBPD
FOXM1
STAT3
EP300
100 200 300
Combined Score
p-value
2e-9
5e-10
Overlap
0.04
0.06
0.08
NPPA/NPPB
ZNF217
WT1
AR
STAT3
SMAD4
PPAR
PAX3-FKHR
SMARCD1
ESR1
CTNNB1
MEF2A
PIAS1
TEAD4
CEBPD
SOX9
SMARCA4
TBX20
TBX20
YAP1
CEBPD
TP53
TCF4
EP300
FOXA2
SMAD3
100 150 200
Combined Score
p-value
2e-8
5e-9
Overlap
0.040
0.045
0.050
0.055
0.060
0.065

**Figure S10. Supplement to Figure 3 – Cardiomyocytes**

**A**, Z-score feature plots for transcriptional signatures enriched in each cardiomyocyte state. Genes used for cell type identification (blue) were selected based on enrichment from Seurat differential expression analysis. **B,** WikiPathways analysis identifies pathways differentially enriched by disease state. Genes used in the analysis included the intersection of pseudobulk and Seurat differential expression analyses with adjusted $p<0.05$. **C.** Palantir pseudotime trajectory analysis of cardiomyocytes. Entropy vs pseudotime plots of donor and DCM cardiomyocytes colored by disease state (top) and cardiomyocyte state (bottom). **D.** enrichPathway analysis comparing enrichment of pathways between cell states. Genes used in the analysis selected from Seurat differential expression with $p<0.05$ and $log2FC>0.1$. **E.** Transcription factor analysis for ADGRL3 (top) and NPPA/NPPB (bottom) states using ChEA 2016 database (https://maayanlab.cloud/Enrichr). Genes used in the analysis selected from Seurat differential expression with $p<0.05$ and $log2FC>0.1$.

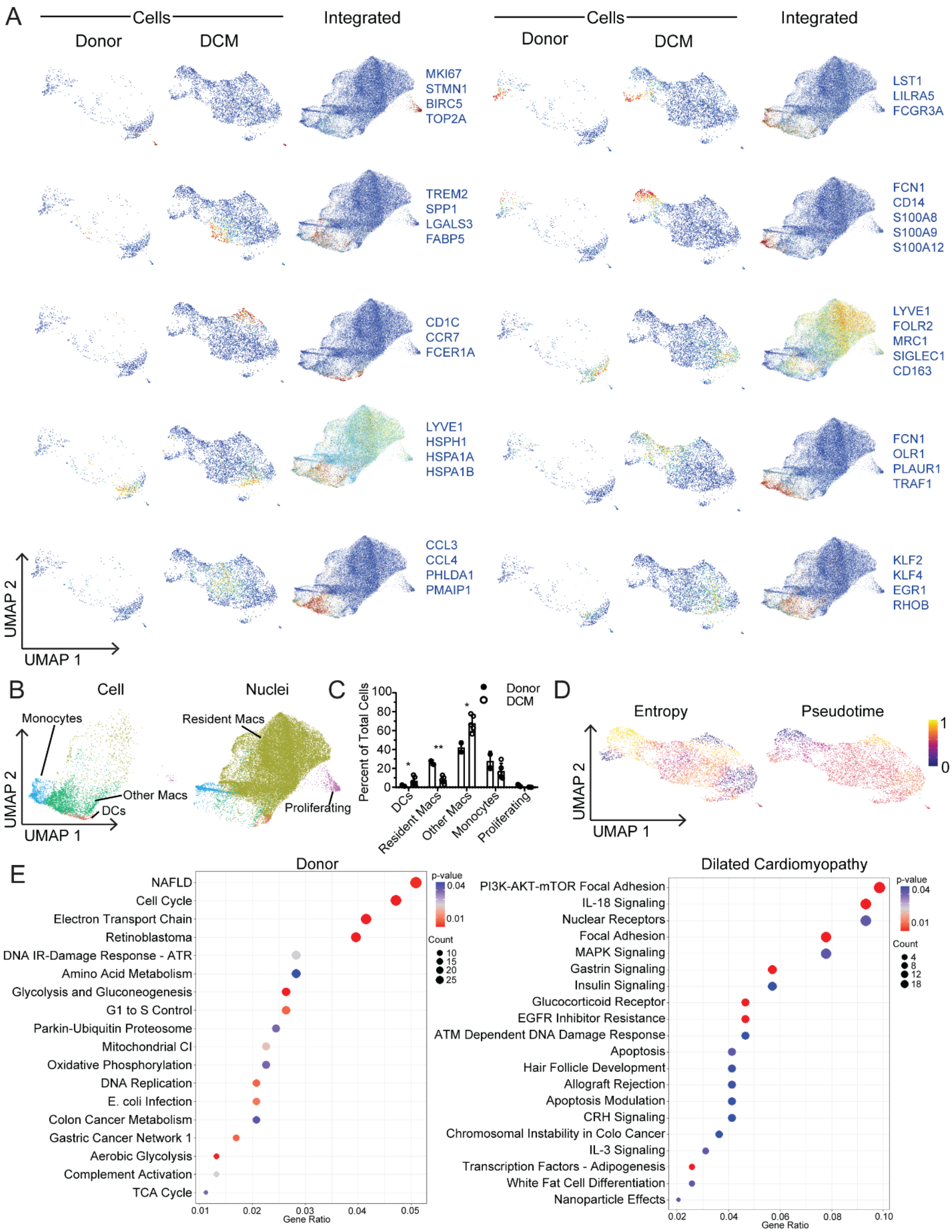
A
Cells
Donor
DCM
Integrated
MKI67
STMN1
BIRC5
TOP2A
LST1
LILRA5
FCGR3A
TREM2
SPP1
LGALS3
FABP5
FCN1
CD14
S100A8
S100A9
S100A12
CD1C
CCR7
FCER1A
LYVE1
FOLR2
MRC1
SIGLEC1
CD163
LYVE1
HSPH1
HSPA1A
HSPA1B
FCN1
OLR1
PLAUR1
TRAF1
CCL3
CCL4
PHLDA1
PMAIP1
KLF2
KLF4
EGR1
RHOB
UMAP 2
UMAP 1
B
Cell
Nuclei
Monocytes
Resident Macs
Other Macs
DCs
Proliferating
C
Percent of Total Cells
Donor
DCM
DCs
Resident Macs
Other Macs
Monocytes
Proliferating
D
Entropy
Pseudotime
E
Donor
NAFLD
Cell Cycle
Electron Transport Chain
Retinoblastoma
DNA IR-Damage Response - ATR
Amino Acid Metabolism
Glycolysis and Gluconeogenesis
G1 to S Control
Parkin-Ubiquitin Proteosome
Mitochondrial CI
Oxidative Phosphorylation
DNA Replication
E. coli Infection
Colon Cancer Metabolism
Gastric Cancer Network 1
Aerobic Glycolysis
Complement Activation
TCA Cycle
Gene Ratio
p-value
Count
Dilated Cardiomyopathy
PI3K-AKT-mTOR Focal Adhesion
IL-18 Signaling
Nuclear Receptors
Focal Adhesion
MAPK Signaling
Gastrin Signaling
Insulin Signaling
Glucocorticoid Receptor
EGFR Inhibitor Resistance
ATM Dependent DNA Damage Response
Apoptosis
Hair Follicle Development
Allograft Rejection
Apoptosis Modulation
CRH Signaling
Chromosomal Instability in Colo Cancer
IL-3 Signaling
Transcription Factors - Adipogenesis
White Fat Cell Differentiation
Nanoparticle Effects
Gene Ratio
p-value
Count

**Figure S11. Supplement to Figure 4 – Monocytes, macrophages, and dendritic cells**

**A**, Z-score feature plots for transcriptional signatures enriched in each monocytes, macrophages, and dendritic cells state. Genes used for cell type identification (blue) were selected based on enrichment from Seurat differential expression analysis. Z-scores are overlaid on the single cell RNA sequencing and integrated UMAP projections. **B.** UMAP projection of the integrated dataset split by technology (single cell vs. single nucleus RNA sequencing) and colored by subpopulation. **C.** Distribution of major cell types recovered from the single cell RNA sequencing dataset (*<0.05, **<0.01, ***<0.001 by Welch's T-test, two-tailed, lines represent mean and standard deviation). **D.** Palantir pseudotime trajectory analysis of monocytes, macrophages, and dendritic cells within the single cell RNA sequencing dataset. Entropy (left) and pseudotime (right) values are overlaid on the UMAP projection. **E,** WikiPathways analysis identifies pathways differentially enriched by disease state. Genes used in the analysis included the intersection of pseudobulk and Seurat differential expression analyses with adjusted p<0.05.

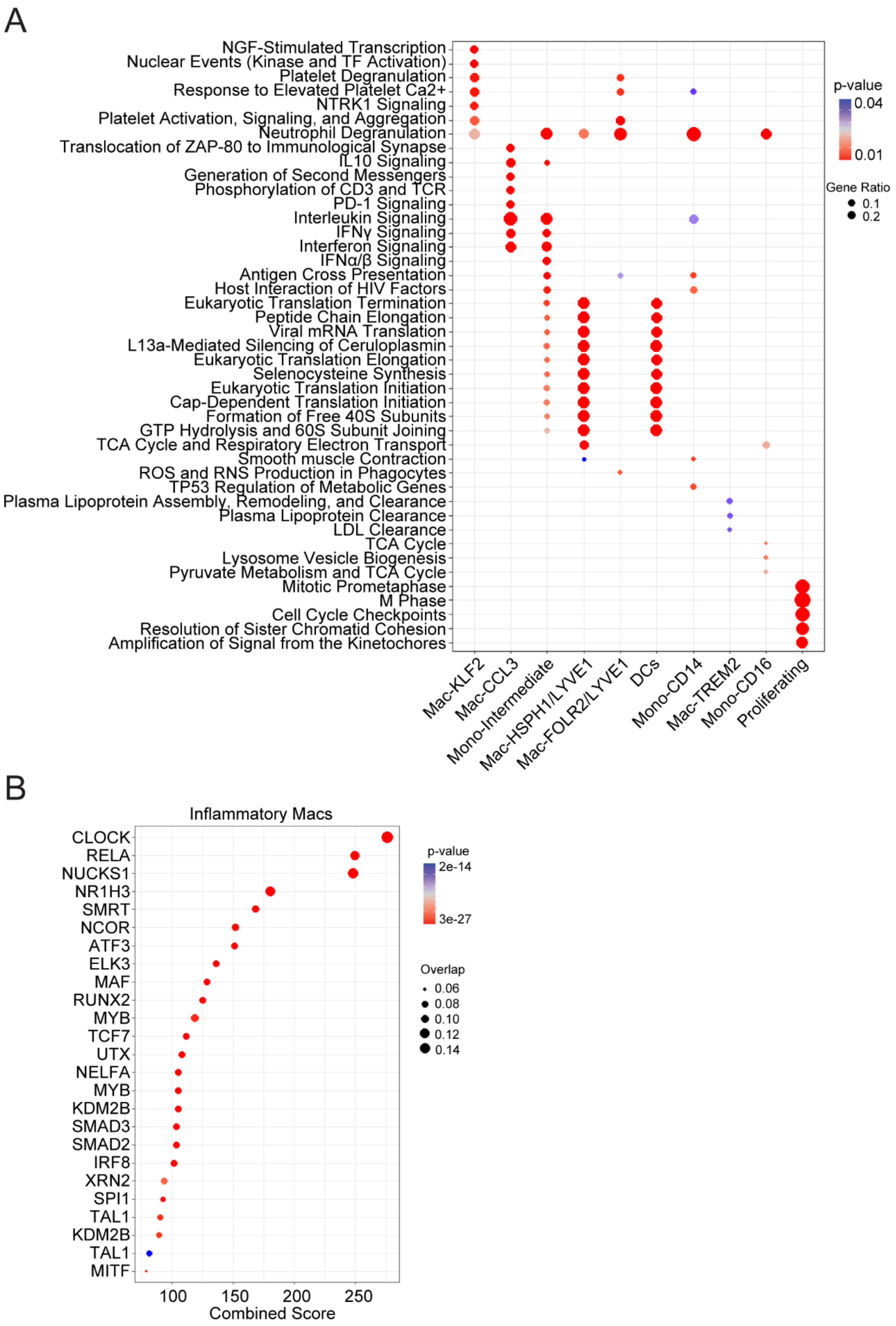
A
NGF-Stimulated Transcription
Nuclear Events (Kinase and TF Activation)
Platelet Degranulation
Response to Elevated Platelet Ca2+
NTRK1 Signaling
Platelet Activation, Signaling, and Aggregation
Neutrophil Degranulation
Translocation of ZAP-80 to Immunological Synapse
IL10 Signaling
Generation of Second Messengers
Phosphorylation of CD3 and TCR
PD-1 Signaling
Interleukin Signaling
IFNγ Signaling
Interferon Signaling
IFNα/β Signaling
Antigen Cross Presentation
Host Interaction of HIV Factors
Eukaryotic Translation Termination
Peptide Chain Elongation
Viral mRNA Translation
L13a-Mediated Silencing of Ceruloplasmin
Eukaryotic Translation Elongation
Selenocysteine Synthesis
Eukaryotic Translation Initiation
Cap-Dependent Translation Initiation
Formation of Free 40S Subunits
GTP Hydrolysis and 60S Subunit Joining
TCA Cycle and Respiratory Electron Transport
Smooth muscle Contraction
ROS and RNS Production in Phagocytes
TP53 Regulation of Metabolic Genes
Plasma Lipoprotein Assembly, Remodeling, and Clearance
Plasma Lipoprotein Clearance
LDL Clearance
TCA Cycle
Lysosome Vesicle Biogenesis
Pyruvate Metabolism and TCA Cycle
Mitotic Prometaphase
M Phase
Cell Cycle Checkpoints
Resolution of Sister Chromatid Cohesion
Amplification of Signal from the Kinetochores
Mac-KLF2
Mac-CCL3
Mono-Intermediate
Mac-HSPH1/LYVE1
Mac-FOLR2/LYVE1
DCs
Mono-CD14
Mac-TREM2
Mono-CD16
Proliferating
p-value
0.04
0.01
Gene Ratio
0.1
0.2
B
Inflammatory Macs
CLOCK
RELA
NUCKS1
NR1H3
SMRT
NCOR
ATF3
ELK3
MAF
RUNX2
MYB
TCF7
UTX
NELFA
MYB
KDM2B
SMAD3
SMAD2
IRF8
XRN2
SPI1
TAL1
KDM2B
TAL1
MITF
100
150
200
250
Combined Score
p-value
2e-14
3e-27
Overlap
0.06
0.08
0.10
0.12
0.14

**Figure S12. Additional Supplement to Figure 4 – Monocytes, macrophages, and dendritic cells**

**A.** enrichPathway analysis comparing enrichment of pathways between cell states. Genes used in the analysis selected from Seurat differential expression with $p<0.05$ and log2FC>0.1. **B.** Transcription factor analysis for genes upregulated in inflammatory macrophage states (KLF2, CCL3, and TREM2) using ChEA 2016 database (https://maayanlab.cloud/Enrichr). Genes used in the analysis selected from Seurat differential expression with $p<0.05$ and log2FC>0.1.

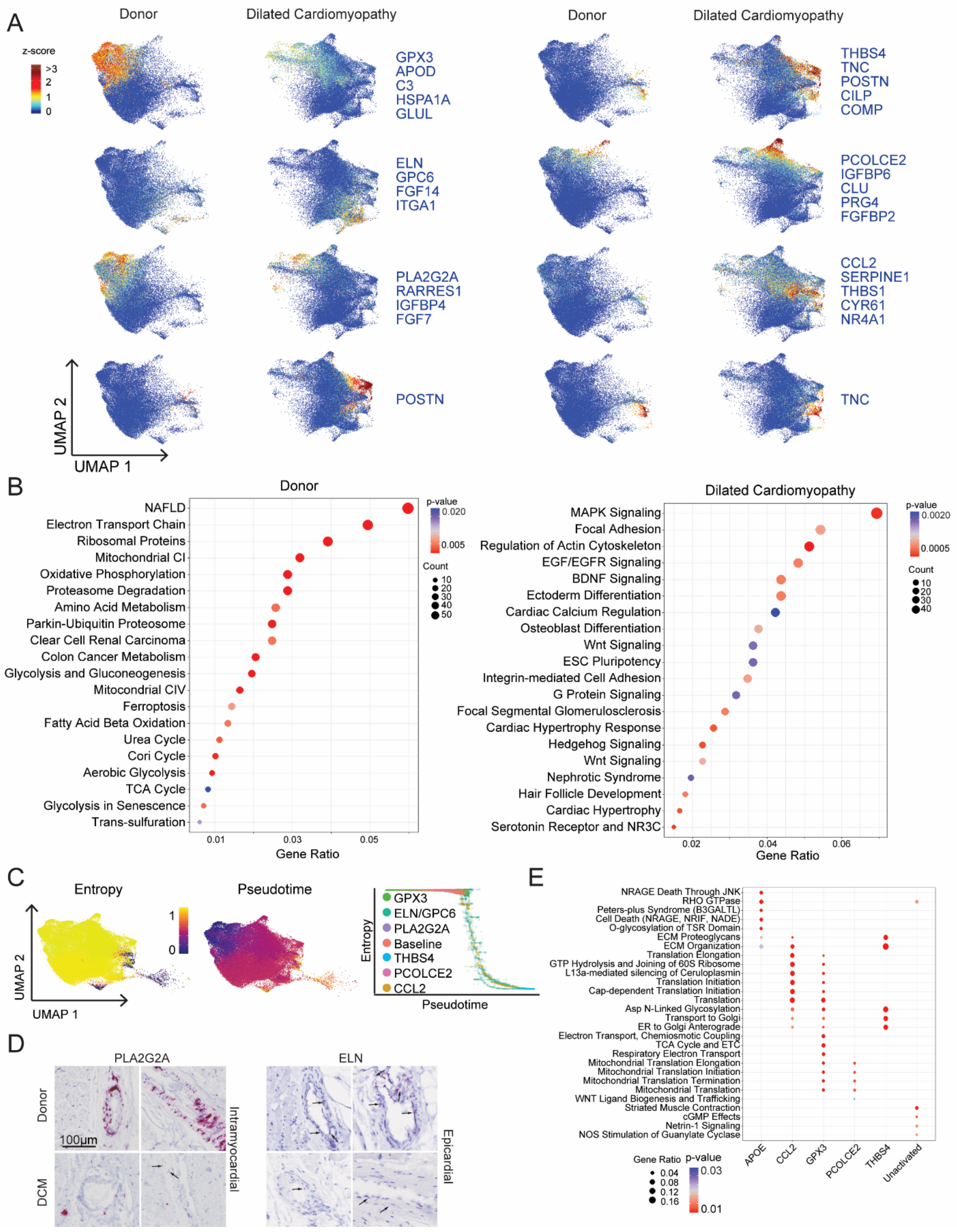
A
Donor
Dilated Cardiomyopathy
Donor
Dilated Cardiomyopathy
z-score
>3
2
1
0
GPX3
APOD
C3
HSPA1A
GLUL
THBS4
TNC
POSTN
CILP
COMP
ELN
GPC6
FGF14
ITGA1
PCOLCE2
IGFBP6
CLU
PRG4
FGFBP2
PLA2G2A
RARRES1
IGFBP4
FGF7
CCL2
SERPINE1
THBS1
CYR61
NR4A1
POSTN
TNC
UMAP 2
UMAP 1
B
Donor
NAFLD
Electron Transport Chain
Ribosomal Proteins
Mitochondrial CI
Oxidative Phosphorylation
Proteasome Degradation
Amino Acid Metabolism
Parkin-Ubiquitin Proteosome
Clear Cell Renal Carcinoma
Colon Cancer Metabolism
Glycolysis and Gluconeogenesis
Mitochondrial CIV
Ferroptosis
Fatty Acid Beta Oxidation
Urea Cycle
Cori Cycle
Aerobic Glycolysis
TCA Cycle
Glycolysis in Senescence
Trans-sulfuration
0.01
0.03
0.05
Gene Ratio
p-value
0.020
0.005
Count
10
20
30
40
50
Dilated Cardiomyopathy
MAPK Signaling
Focal Adhesion
Regulation of Actin Cytoskeleton
EGF/EGFR Signaling
BDNF Signaling
Ectoderm Differentiation
Cardiac Calcium Regulation
Osteoblast Differentiation
Wnt Signaling
ESC Pluripotency
Integrin-mediated Cell Adhesion
G Protein Signaling
Focal Segmental Glomerulosclerosis
Cardiac Hypertrophy Response
Hedgehog Signaling
Wnt Signaling
Nephrotic Syndrome
Hair Follicle Development
Cardiac Hypertrophy
Serotonin Receptor and NR3C
0.02
0.04
0.06
Gene Ratio
p-value
0.0020
0.0005
Count
10
20
30
40
C
Entropy
Pseudotime
1
0
UMAP 2
UMAP 1
GPX3
ELN/GPC6
PLA2G2A
Baseline
THBS4
PCOLCE2
CCL2
Entropy
Pseudotime
D
PLA2G2A
ELN
Donor
DCM
100µm
Intramyocardial
Epicardial
E
NRAGE Death Through JNK
RHO GTPase
Peters-plus Syndrome (B3GALTL)
Cell Death (NRAGE, NRIF, NADE)
O-glycosylation of TSR Domain
ECM Proteoglycans
ECM Organization
Translation Elongation
GTP Hydrolysis and Joining of 60S Ribosome
L13a-mediated silencing of Ceruloplasmin
Translation Initiation
Cap-dependent Translation Initiation
Translation
Asp N-Linked Glycosylation
Transport to Golgi
ER to Golgi Anterograde
Electron Transport, Chemiosmotic Coupling
TCA Cycle and ETC
Respiratory Electron Transport
Mitochondrial Translation Elongation
Mitochondrial Translation Initiation
Mitochondrial Translation Termination
Mitochondrial Translation
WNT Ligand Biogenesis and Trafficking
Striated Muscle Contraction
cGMP Effects
Netrin-1 Signaling
NOS Stimulation of Guanylate Cyclase
APOE
CCL2
GPX3
PCOLCE2
THBS4
Unactivated
Gene Ratio
0.04
0.08
0.12
0.16
p-value
0.03
0.01

**Figure S13. Supplement to Figure 5 – Fibroblasts**

**A**, Z-score feature plots for transcriptional signatures enriched in each fibroblast state. Genes used for cell type identification (blue) were selected based on enrichment from Seurat differential expression analysis. Z-scores are overlaid on the integrated UMAP projections split by disease state. **B,** WikiPathways analysis identifies pathways differentially enriched by disease state. Genes used in the analysis included the intersection of pseudobulk and Seurat differential expression analyses with adjusted $p<0.05$. **C,** Palantir pseudotime trajectory analysis of fibroblasts. Entropy and pseudotime values are overlaid on the UMAP projection (left). Entropy vs pseudotime plots identifying differing fibroblast trajectories (right). Fibroblast expressing CCL2, THBS4, and ELN were identified as the most differentiated cell states. **D,** RNA *in situ* hybridization for PLA2G2A and ELN (red). Representative images showing perivascular staining of PLA2G2A in the myocardium of donor samples. Minimal staining was observed in DCM samples. ELN staining was observed in the media of epicardial coronary arteries in both donor and DCM samples. **E.** enrichPathway analysis comparing enrichment of pathways between cell states. Genes used in the analysis selected from Seurat differential expression with $p<0.05$ and $log2FC>0.1$.

A

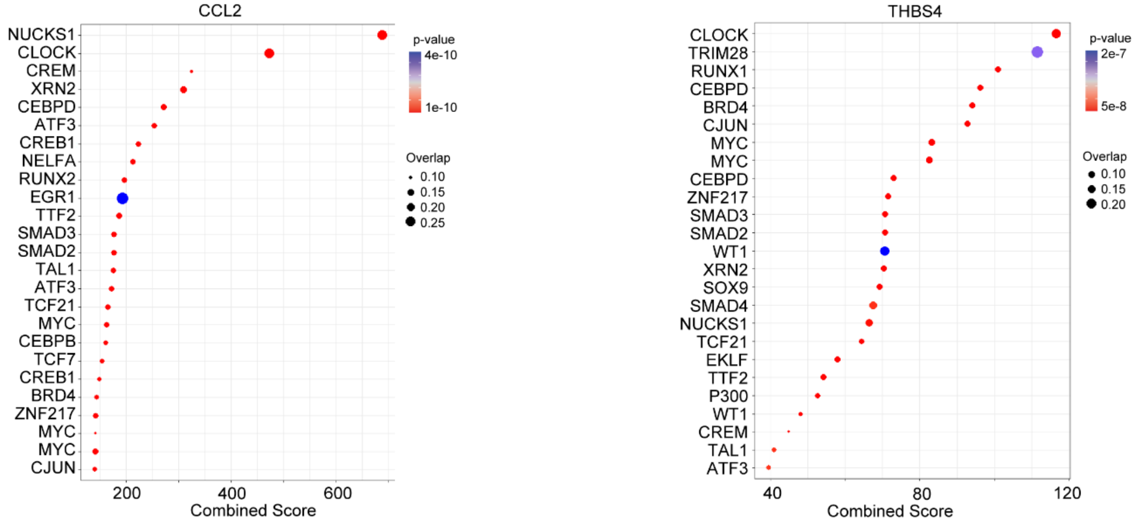

CCL2
NUCKS1
CLOCK
CREM
XRN2
CEBPD
ATF3
CREB1
NELFA
RUNX2
EGR1
TTF2
SMAD3
SMAD2
TAL1
ATF3
TCF21
MYC
CEBPB
TCF7
CREB1
BRD4
ZNF217
MYC
MYC
CJUN
200
400
600
Combined Score
p-value
4e-10
1e-10
Overlap
0.10
0.15
0.20
0.25
THBS4
CLOCK
TRIM28
RUNX1
CEBPD
BRD4
CJUN
MYC
MYC
CEBPD
ZNF217
SMAD3
SMAD2
WT1
XRN2
SOX9
SMAD4
NUCKS1
TCF21
EKLF
TTF2
P300
WT1
CREM
TAL1
ATF3
40
80
120
Combined Score
p-value
2e-7
5e-8
Overlap
0.10
0.15
0.20

**Figure S14. Additional Supplement to Figure 5 – Fibroblasts**

**A.** Transcription factor analysis for CCL2 (left) and THBS4 (right) states using ChEA 2016 database (https://maayanlab.cloud/Enrichr). Genes used in the analysis selected from Seurat differential expression with $p<0.05$ and $\log2FC>0.1$.

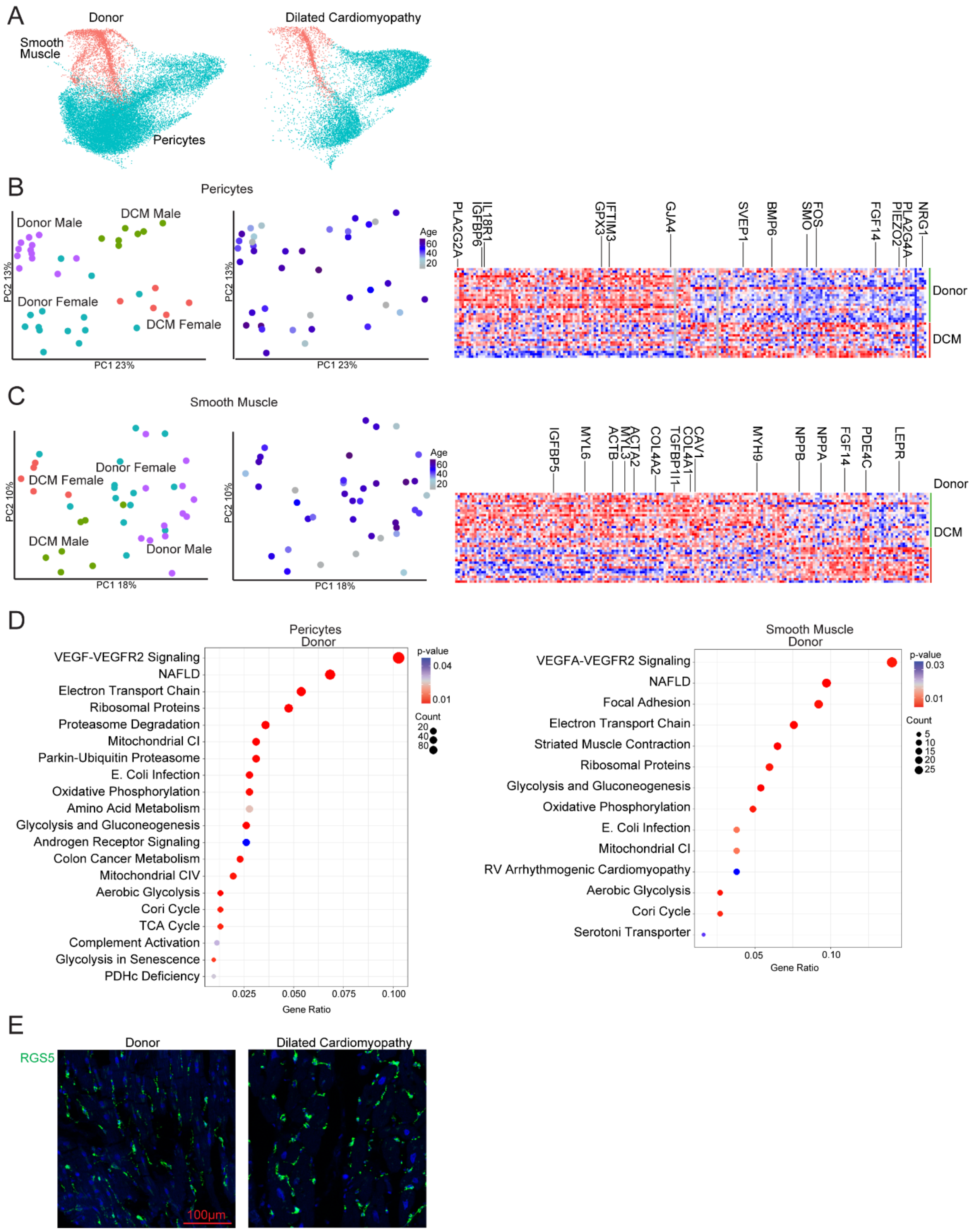
A
Donor
Dilated Cardiomyopathy
Smooth Muscle
Pericytes
B
Pericytes
Donor Male
DCM Male
Donor Female
DCM Female
PC1 23%
PC2 13%
Age
60
40
20
PLA2G2A
IGFBP6
IL18R1
GPX3
IFTIM3
GJA4
SVEP1
BMP6
SMO
FOS
FGF14
PIEZO2
PLA2G4A
NRG1
Donor
DCM
C
Smooth Muscle
Donor Female
DCM Female
DCM Male
Donor Male
PC1 18%
PC2 10%
IGFBP5
MYL6
ACTB
MYL3
ACTA2
COL4A2
TGFBP1I1
COL4A1
CAV1
MYH9
NPPB
NPPA
FGF14
PDE4C
LEPR
D
Pericytes
Donor
VEGF-VEGFR2 Signaling
NAFLD
Electron Transport Chain
Ribosomal Proteins
Proteasome Degradation
Mitochondrial CI
Parkin-Ubiquitin Proteasome
E. Coli Infection
Oxidative Phosphorylation
Amino Acid Metabolism
Glycolysis and Gluconeogenesis
Androgen Receptor Signaling
Colon Cancer Metabolism
Mitochondrial CIV
Aerobic Glycolysis
Cori Cycle
TCA Cycle
Complement Activation
Glycolysis in Senescence
PDHc Deficiency
0.025
0.050
0.075
0.100
Gene Ratio
p-value
0.04
0.01
Count
20
40
80
Smooth Muscle
Donor
VEGFA-VEGFR2 Signaling
NAFLD
Focal Adhesion
Electron Transport Chain
Striated Muscle Contraction
Ribosomal Proteins
Glycolysis and Gluconeogenesis
Oxidative Phosphorylation
E. Coli Infection
Mitochondrial CI
RV Arrhythmogenic Cardiomyopathy
Aerobic Glycolysis
Cori Cycle
Serotoni Transporter
0.05
0.10
Gene Ratio
p-value
0.03
0.01
Count
5
10
15
20
25
E
Donor
Dilated Cardiomyopathy
RGS5
100µm

**Figure S15. Pericytes and smooth muscle cells exhibit global changes in gene expression in dilated cardiomyopathy.**

**A,** Unsupervised clustering of pericytes and fibroblasts within the integrated dataset split by disease state. **B-C,** Principal component analysis (PCA, DESeq2) plots of pericyte (B) and smooth muscle cell (C) pseudobulk single nucleus RNA sequencing data colored by sex and disease state and age. Each data point represents an individual subject. Heatmaps displaying the top 100 upregulated and downregulated genes ranked by log2 fold-change comparing donor control to dilated cardiomyopathy (DCM). Differentially expressed genes were derived from the intersection of pseudobulk (DESeq2) and single cell (Seurat) analyses. **D,** WikiPathways analysis identifies pathways differentially enriched in pericytes (left) and smooth muscle cells (right) by disease state. Genes used in the analysis included the intersection of pseudobulk and Seurat differential expression analyses with adjusted $p<0.05$. **E.** Representative images of RGS5 staining for pericytes by RNA *in situ* hybridization.

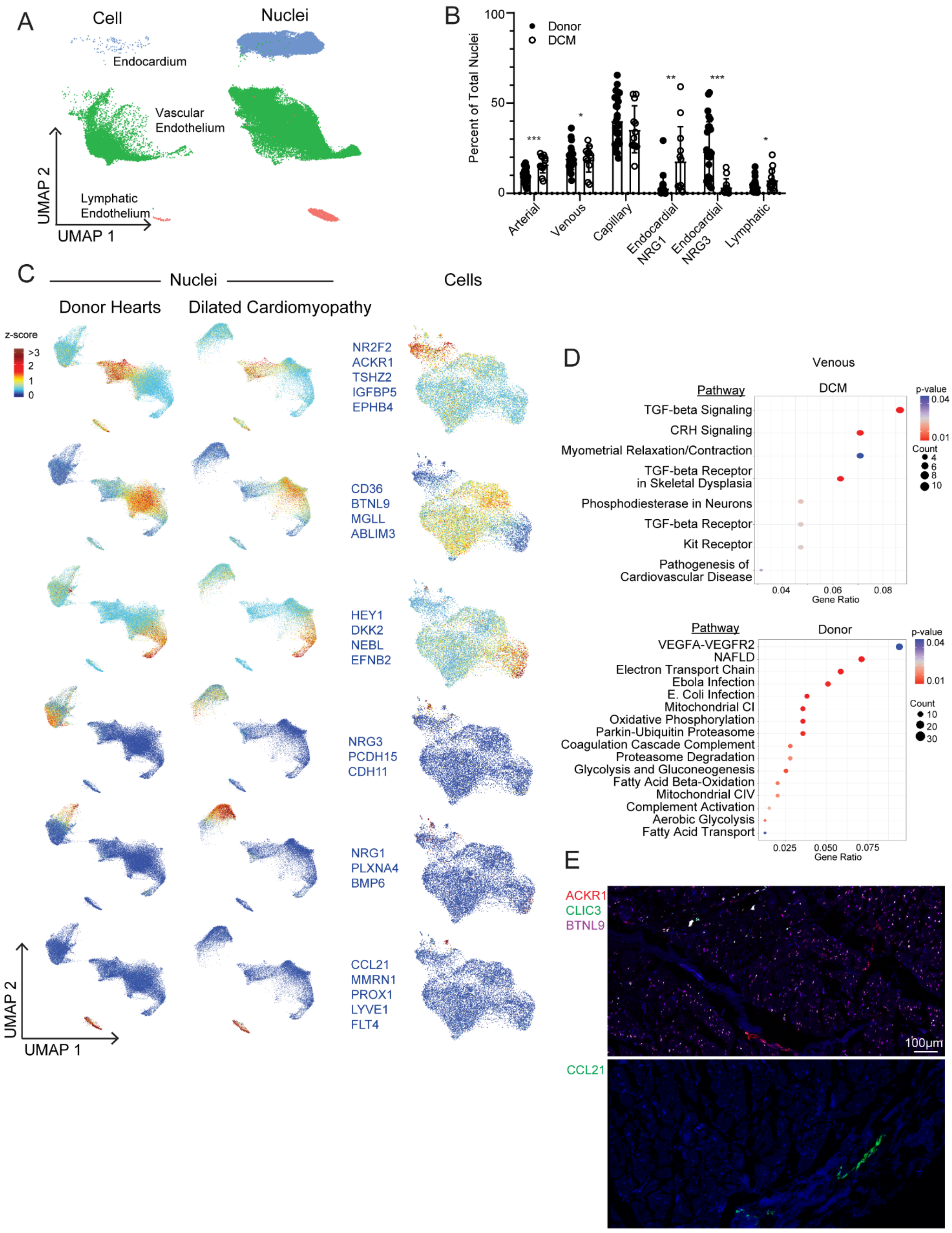
A
Cell
Nuclei
Endocardium
Vascular
Endothelium
Lymphatic
Endothelium
UMAP 2
UMAP 1
B
100
50
0
Percent of Total Nuclei
Donor
DCM
***
*
**
***
*
Arterial
Venous
Capillary
Endocardial NRG1
Endocardial NRG3
Lymphatic
C
Nuclei
Cells
Donor Hearts
Dilated Cardiomyopathy
z-score
>3
2
1
0
NR2F2
ACKR1
TSHZ2
IGFBP5
EPHB4
CD36
BTNL9
MGLL
ABLIM3
HEY1
DKK2
NEBL
EFNB2
NRG3
PCDH15
CDH11
NRG1
PLXNA4
BMP6
CCL21
MMRN1
PROX1
LYVE1
FLT4
UMAP 2
UMAP 1
D
Venous
Pathway
DCM
p-value
0.04
0.01
Count
4
6
8
10
TGF-beta Signaling
CRH Signaling
Myometrial Relaxation/Contraction
TGF-beta Receptor
in Skeletal Dysplasia
Phosphodiesterase in Neurons
TGF-beta Receptor
Kit Receptor
Pathogenesis of
Cardiovascular Disease
0.04
0.06
0.08
Gene Ratio
Pathway
Donor
p-value
0.04
0.01
Count
10
20
30
VEGFA-VEGFR2
NAFLD
Electron Transport Chain
Ebola Infection
E. Coli Infection
Mitochondrial CI
Oxidative Phosphorylation
Parkin-Ubiquitin Proteasome
Coagulation Cascade Complement
Proteasome Degradation
Glycolysis and Gluconeogenesis
Fatty Acid Beta-Oxidation
Mitochondrial CIV
Complement Activation
Aerobic Glycolysis
Fatty Acid Transport
0.025
0.050
0.075
Gene Ratio
E
ACKR1
CLIC3
BTNL9
100µm
CCL21

**Figure S16. Supplement to Figure 6 – Endothelial cells**

**A,** UMAP projection of the integrated dataset split by technology (single cell vs. single nucleus RNA sequencing) and colored by subpopulation. **B,** Distribution of nuclei in the integrated object divided by major cell type (*<0.05, **<0.01, ***<0.001 by Welch's T-test, two-tailed, lines represent mean and standard deviation). **C,** Z-score feature plots for transcriptional signatures enriched in endothelial and endocardial cell populations. Genes used for cell type identification (blue) were selected based on enrichment from Seurat differential expression analysis. Z-scores are overlaid on the single nucleus RNA sequencing (split by disease) and single cell RNA sequencing UMAP projections. **D,** WikiPathways analysis identifies pathways differentially enriched in venous endothelial cells by disease state. Genes used in the analysis included the intersection of pseudobulk and Seurat differential expression analyses with adjusted p<0.05. **E.** Representative RNAScope images of vascular (top) and lymphatic (bottom) endothelial cells. ACKR1 – venous, CLIC3 – arterial, BTNL9 – capillary, CCL21 – lymphatic.

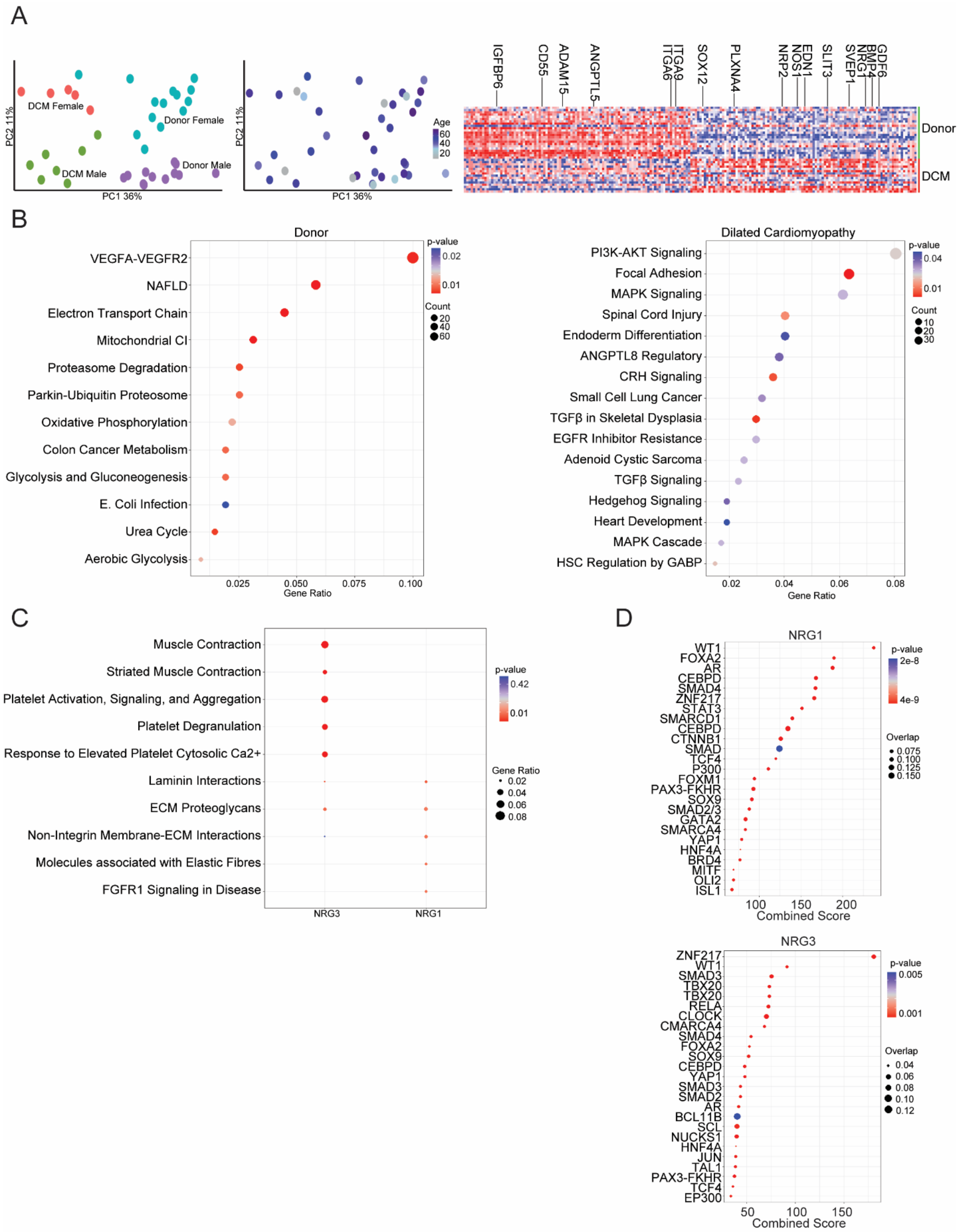
A
DCM Female
Donor Female
DCM Male
Donor Male
PC2 11%
PC1 36%
Age
60
40
20
IGFBP6
CD55
ADAM15
ANGPTL5
ITGA6
ITGA9
SOX12
PLXNA4
NRP2
NOS1
EDN1
SLIT3
SVEP1
NRG1
BMP4
GDF6
Donor
DCM
B
Donor
VEGFA-VEGFR2
NAFLD
Electron Transport Chain
Mitochondrial CI
Proteasome Degradation
Parkin-Ubiquitin Proteosome
Oxidative Phosphorylation
Colon Cancer Metabolism
Glycolysis and Gluconeogenesis
E. Coli Infection
Urea Cycle
Aerobic Glycolysis
Gene Ratio
p-value
Count
Dilated Cardiomyopathy
PI3K-AKT Signaling
Focal Adhesion
MAPK Signaling
Spinal Cord Injury
Endoderm Differentiation
ANGPTL8 Regulatory
CRH Signaling
Small Cell Lung Cancer
TGFβ in Skeletal Dysplasia
EGFR Inhibitor Resistance
Adenoid Cystic Sarcoma
TGFβ Signaling
Hedgehog Signaling
Heart Development
MAPK Cascade
HSC Regulation by GABP
Gene Ratio
C
Muscle Contraction
Striated Muscle Contraction
Platelet Activation, Signaling, and Aggregation
Platelet Degranulation
Response to Elevated Platelet Cytosolic Ca2+
Laminin Interactions
ECM Proteoglycans
Non-Integrin Membrane-ECM Interactions
Molecules associated with Elastic Fibres
FGFR1 Signaling in Disease
NRG3
NRG1
p-value
Gene Ratio
D
NRG1
WT1
FOXA2
AR
CEBPD
SMAD4
ZNF217
STAT3
SMARCD1
CEBPD
CTNNB1
SMAD
TCF4
P300
FOXM1
PAX3-FKHR
SOX9
SMAD2/3
GATA2
SMARCA4
YAP1
HNF4A
BRD4
MITF
OLI2
ISL1
Combined Score
Overlap
NRG3
ZNF217
WT1
SMAD3
TBX20
TBX20
RELA
CLOCK
CMARCA4
SMAD4
FOXA2
SOX9
CEBPD
YAP1
SMAD3
SMAD2
AR
BCL11B
SCL
NUCKS1
HNF4A
JUN
TAL1
PAX3-FKHR
TCF4
EP300
Combined Score

**Figure S17. Supplement to endocardial panels of Figure 6**

**A,** Principal component analysis (PCA, DESeq2) plots of endocardial cell pseudobulk single nucleus RNA sequencing data colored by sex and disease state and age. Each data point represents an individual subject. Heatmaps displaying the top 100 upregulated and downregulated genes ranked by log2 fold-change comparing donor control to dilated cardiomyopathy (DCM). Differentially expressed genes were derived from the intersection of pseudobulk (DESeq2) and single cell (Seurat) analyses. **B,** WikiPathways analysis identifies pathways differentially enriched in endocardial cells by disease state. Genes used in the analysis included the intersection of pseudobulk and Seurat differential expression analyses with adjusted $p<0.05$. **C.** enrichPathway analysis comparing enrichment of pathways between cell states. Genes used in the analysis selected from Seurat differential expression with $p<0.05$ and $\log2FC>0.1$. **D.** Transcription factor analysis for NRG1 (top) and NRG3 (bottom) states using ChEA 2016 database (https://maayanlab.cloud/Enrichr). Genes used in the analysis selected from Seurat differential expression with $p<0.05$ and $\log2FC>0.1$.

| | |
|---|---|
| **BIOSPECIMEN TYPE** | Solid Tissue |
| **ANATOMICAL SITE** | Heart, Left-ventricle myocardium |
| **DISEASE STATUS OF PATIENTS** | Heart failure, Healthy controls |
| **CLINICAL CHARACTERISTICS OF PATIENTS** | Dilated (Non-ischemic) cardiomyopathy, Healthy donors |
| **VITAL STATE OF PATIENTS** | Alive with LVAD or transplant, Deceased |
| **COLLECTION MECHANISM** | LVAD coring device, Scalpel dissection |
| **TYPE OF STABILIZATION** | On ice |
| **TYPE OF LONG-TERM PRESERVATION** | Flash frozen in liquid nitrogen |
| **STORAGE TERMPATURE** | -80°C |
| **STORAGE DURATION** | 1 day - 9 years |
| **COMPOSITION ASSESSMENT & SELECTION** | Myocardium tissue sufficient to yield 100,000+ cells/nuclei per sample after digest and dissociation |
| **YEAR SAMPLES PROCESSED** | 2019 |

**Table S4. BRISQ Data for Human Specimens.**
